# A host-directed adjuvant resuscitates and sensitizes intracellular bacterial persisters to antibiotics

**DOI:** 10.1101/2024.09.30.615828

**Authors:** Kuan-Yi Lu, Xiangbo Yang, Matthew J.G. Eldridge, Nikki J. Wagner, Brian Hardy, Matthew Axtman, Sarah E. Rowe, Xiaodong Wang, Vance G. Fowler, Sophie Helaine, Kenneth H. Pearce, Brian P. Conlon

## Abstract

There are two major problems in the field of antimicrobial chemotherapy–antibiotic resistance and antibiotic tolerance. In the case of antibiotic tolerance, antibiotics fail to kill the bacteria as their phenotypic state affords them protection from the bactericidal activity of the antibiotic. Antibiotic tolerance can affect an entire bacterial population, or a subset of cells known as persister cells. Interaction with the host induces the formation of persister cells in numerous pathogens, with reactive oxygen and nitrogen species production being heavily implicated in the collapse of bacterial energy levels and entrance into an antibiotic tolerant state. Here, we developed a high-throughput screen to identify energy modulators for intracellular *Staphylococcus aureus*. The identified compound, **KL1**, increases intracellular bacterial energy and sensitizes the persister population to antibiotics, without causing cytotoxicity or bacterial outgrowth. We demonstrate that **KL1** exhibits adjuvant activity in a murine model of *S. aureus* bacteremia and intracellular infection of *Salmonella Typhimurium*. Transcriptomic analysis and further studies on its mechanism of action reveal that **KL1** modulates host immune response genes and suppresses the production of reactive species in host macrophages, alleviating one of the major stressors that induce antibiotic tolerance. Our findings highlight the potential to target intracellular persister cells by stimulating their metabolism and encourage larger efforts to address antibiotic tolerance at the host–pathogen interface, particularly within the intracellular milieu.

## Introduction

Antibiotic tolerance has been frequently connected with poor treatment outcomes in the clinic (1–3). Unlike antibiotic resistance, which permits bacterial growth in the presence of drugs, antibiotic tolerance allows bacteria to withstand multiple antibiotics for prolonged periods (4, 5). This phenomenon is best illustrated by clinical observations in which bacterial isolates that fail to respond to treatment in patients remain susceptible to the prescribed antibiotics by standard minimum inhibitory concentration (MIC) testing (6, 7). The extended survival of tolerant bacteria further predisposes them to evolve antibiotic resistance over time, underscoring the critical need to address antibiotic tolerance (8, 9).

A principal characteristic of tolerant bacteria is their lower energy level and metabolic activities, rendering a non-growing state which prevents them from being targeted by antibiotics that are directed at growth-centered processes (3, 10–14). Antibiotic tolerance can be induced by the infection environment and the activities of innate immune cells. The intracellular environment in macrophages has been strongly associated with the induction of metabolic indolence and antibiotic tolerance (15–19). Importantly, host-produced reactive oxygen and nitrogen species are responsible for the induction of antibiotic tolerance in a wide range of bacterial species, including *Staphylococcus aureus*, *Mycobacterium tuberculosis*, *Yersinia pseudotuberculosis* and *Salmonella Typhimurium* (15, 16, 18–24). Additional intracellular stressors, such as phagosome acidification and nutrient deprivation, can also contribute to antibiotic tolerance (18). When conditions become favorable, the intracellular bacterial persisters can resume growth, leading to relapsing infections. Therefore, developing innovative strategies to target intracellular bacterial persisters is essential for achieving complete eradication.

To address this difficult-to-eradicate intracellular reservoir, we established a high-throughput screening platform to identify potential energy modulators, aiming to sensitize these bacteria to antibiotics. Here, we targeted *S. aureus* as this versatile facultative intracellular pathogen is capable of surviving within various cell types (25–29) and frequently causes treatment failure and recurrent infections due to antibiotic tolerance, despite appropriate use of antibiotics (7, 30). Among the >4,700 drug-like compounds profiled, we identified that the lead compound **KL1** can resuscitate intracellular *S. aureus* and enhance antibiotic efficacy across multiple clinical isolates and laboratory strains, including methicillin-resistant *S. aureus* (MRSA). We confirm that treatment with **KL1** alone does not induce bacterial outgrowth and shows no detectable cytotoxicity in mammalian cells. Additionally, **KL1** exhibits adjuvant activity in a murine model of *S. aureus* bacteremia and intracellular infection of *Salmonella Typhimurium* within macrophages. Using transcriptomic, chemical and biochemical approaches, we discover that, rather than directly targeting the bacteria, **KL1** consistently reduces the levels of reactive oxygen and/or nitrogen species, likely through targeting the upstream regulator EHMT2/G9a. Together, we demonstrate the potential to pharmacologically sensitize intracellular bacteria to antibiotics by altering the host environment. Our energy modulator screening platform provides a pathway to uncover host-directed therapies that synergize with antibiotics and aid in the clearance of intracellular persisters.

## Results

### Intracellular environment is an important niche driving antibiotic tolerant *S. aureus*

Antibiotic tolerance measured in test tubes is often poorly correlated with clinical outcomes in patients, highlighting the importance of probing bacterial drug responses within the context of host–pathogen interactions (6). Multiple lines of evidence suggest that host intracellular environment serves as a privileged site of infection that antagonizes antimicrobial agents for obligate and facultative intracellular bacterial pathogens, including *S. aureus* (15, 17, 31–33). This phenomenon can be illustrated by testing clinical isolates with varying persister frequencies in tube assays with cell-based models. We found that although two *S. aureus* isolates displayed a dramatic 500-fold difference in persister formation in tubes, both produced similar high-level antibiotic tolerance following internalization by bone marrow-derived macrophages (BMDMs) (**Figure 1A** and **B**). Both isolates produced more tolerant bacteria inside the macrophages (46–78% of the intracellular bacterial population) compared to their planktonic cultures (0.03–15% of the extracellular bacteria) (**Figure 1A** and **B**). These isolates had the same minimum inhibitory concentrations (MICs) for rifampicin (6 to 8 ± 4 ng/mL) and other antibiotics used (Supplementary table 2). This data indicates that the host interaction plays a dominant role in promoting antibiotic tolerance.

**Figure 1.**
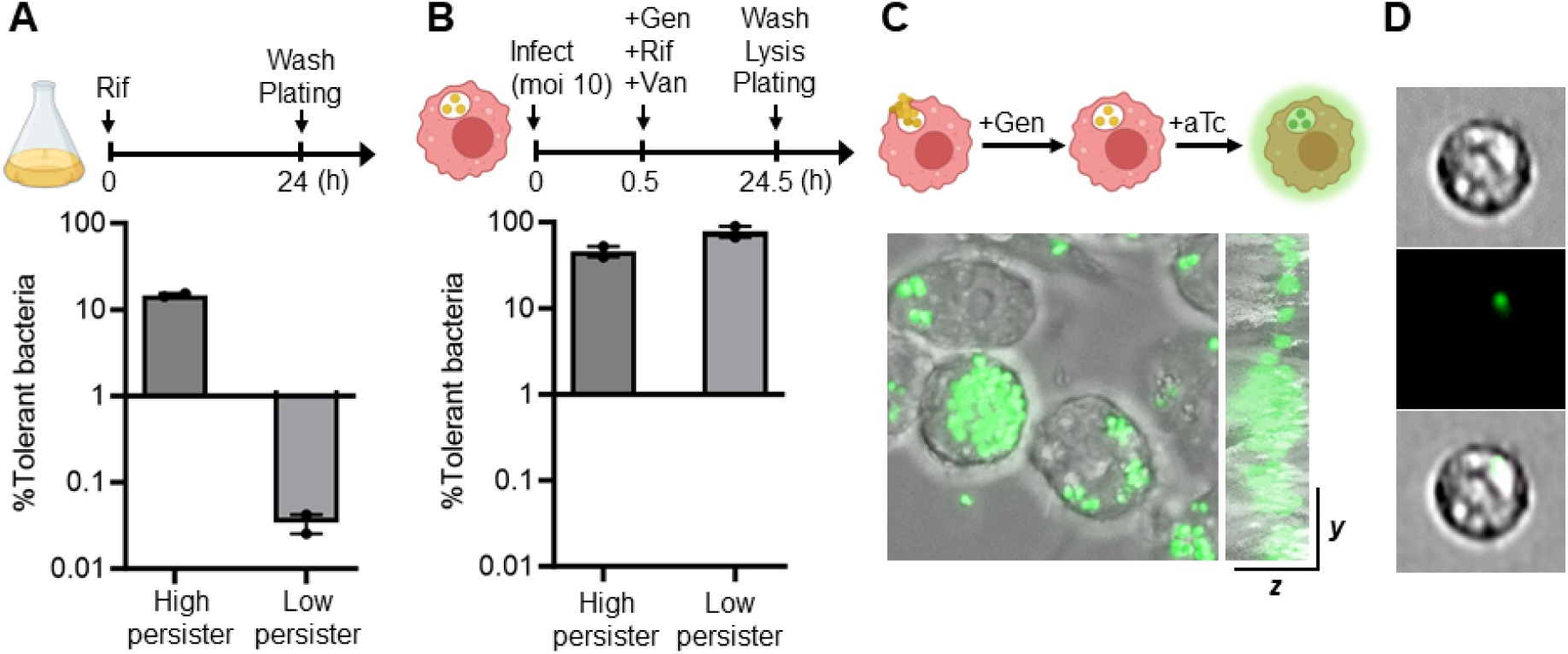
Intracellular environments provide an important niche producing antibiotic-tolerant *S. aureus* persisters. (**A**) Clinical isolates from *S. aureus* bacteremia patients exhibited variable antibiotic tolerability in planktonic cultures. Rifampicin (Rif) was washed away to enumerate the numbers of survivor bacteria (CFU). (**B**) The high and low persister isolates characterized in (**A**) became more tolerant and produced comparable numbers of persisters in bone marrow-derived macrophages. Gentamicin (Gen) and vancomycin (Van) were administered to eradicate the extracellular bacteria. The numbers of survivor bacteria were normalized to the corresponding numbers of intracellular CFUs before the antibiotic treatment (T_0_) to calculate the persister frequencies. Representative data of three independent experiments are shown (n = 3). The bars represent mean ± SEM. Assay schematics are shown above plots. (**C**) Confocal z-sectioning visualized viable intracellular *S. aureus*. RAW 264.7 macrophages were infected with an inducible GFP reporter strain and treated with 50 µg/mL Gen to exclude extracellular *S. aureus*. Live intracellular bacteria can be probed with anhydrotetracycline (aTc) induction. (**D**) Intracellular persisters in mouse kidneys were detected using ImageStream analysis. C57BL/6J mice were systemically infected with the inducible reporter strain via the intravenous route and intraperitoneally treated with 10 mg/kg Rif at 1 day post-infection (dpi) for 24 h. The kidney cells were extracted at 2 dpi and incubated with 2 µM aTc to induce GFP expression. Representative images are shown.

As a versatile pathogen, *S. aureus* can adapt to multiple intracellular niches and colonize a wide range of mammalian cells. To facilitate tracking intracellular persister cells in vivo, we constructed inducible *S. aureus* fluorescent reporter strains and first demonstrated the feasibility of using these reporter strains to probe intracellular *S. aureus*. We showed that viable bacteria inside macrophages can be visualized using confocal microscopy and 3D rendering following induction with anhydrotetracycline (aTc) (**Figure 1C** and Movie S1– 3). To explore whether *S. aureus* survives antibiotic treatment inside mammalian cells, C57BL/6J mice were infected with the reporter strains via tail vein injections (i.v.) and treated with 10 mg/kg rifampicin via the intraperitoneal (i.p.) route at 1 day post-infection (dpi). Rifampicin was chosen as this drug is one of the few antibiotics that can easily penetrate mammalian cells (15). The kidney cells were then extracted at 2 dpi, followed by aTc treatment to induce GFP expression. The induction was performed in the presence of gentamicin to exclude the extracellular bacteria. Using ImageStream analysis, we observed live cells harboring GFP-expressing *S. aureus* that survived rifampicin treatment (**Figure 1D**). Thus, the intracellular environment not only provides a physical barrier against antimicrobial agents but also promotes antibiotic tolerance during the infection. These findings reiterate the need to devise a strategy for eradicating intracellular persister cells, which is particularly crucial given the limited antibiotic options available against intracellular bacteria.

### A high-throughput screen for energy modulators identifies a compound that sensitizes intracellular *S. aureus* to antibiotics

Professional phagocytes, including macrophages, facilitate the clearance of bacteria during bloodstream infections. However, *S. aureus* can survive this hostile environment after engulfment, by adopting a less metabolically active lifestyle (25). This metabolic indolence at a lower energy state favors antibiotic tolerance and persister cell formation as most antibiotics are directed at growth-centered processes. To address this, we developed a high-throughput platform to screen compounds that resuscitate intracellular *S. aureus*, with the goal of sensitizing the intracellular population to antibiotics (**Figure 2A**). Here, we utilized a bioluminescent methicillin-resistant *S. aureus* (MRSA) strain JE2–lux to probe the energy state of the intracellular bacteria (34). To assess its feasibility, RAW 264.7 macrophages were infected with JE2–lux, followed by antibiotic treatment. As shown in **Figure 2B**, rifampicin reduced the bioluminescent signal in the infected cells, while vancomycin, which does not penetrate the mammalian plasma membrane, had no effect on the intracellular *S. aureus* activity. Next, we incorporated a cell viability assay into the workflow to monitor drugs’ cytotoxicity. We verified that the readout reflected the number of live mammalian cells without being interfered by the presence of bacteria (**Figure 2C**). We then evaluated the dynamic range of the assay by measuring bioluminescent signals as a function of rifampicin concentrations. The dose response analysis indicated that a concentration of 2 ng/mL rifampicin resulted in a 50% reduction in the intracellular bacterial activity without compromising the host cell viability (**Figure 2D**). This EC_50_ value (2 ng/mL) aligned with the rifampicin MIC (6 ng/mL) for the strain JE2–lux, indicating the utility of this platform (Supplementary table 2). In comparison, vancomycin did not affect the bioluminescent signal of intracellular *S. aureus* across different concentrations (**Figure 2E**).

**Figure 2.**
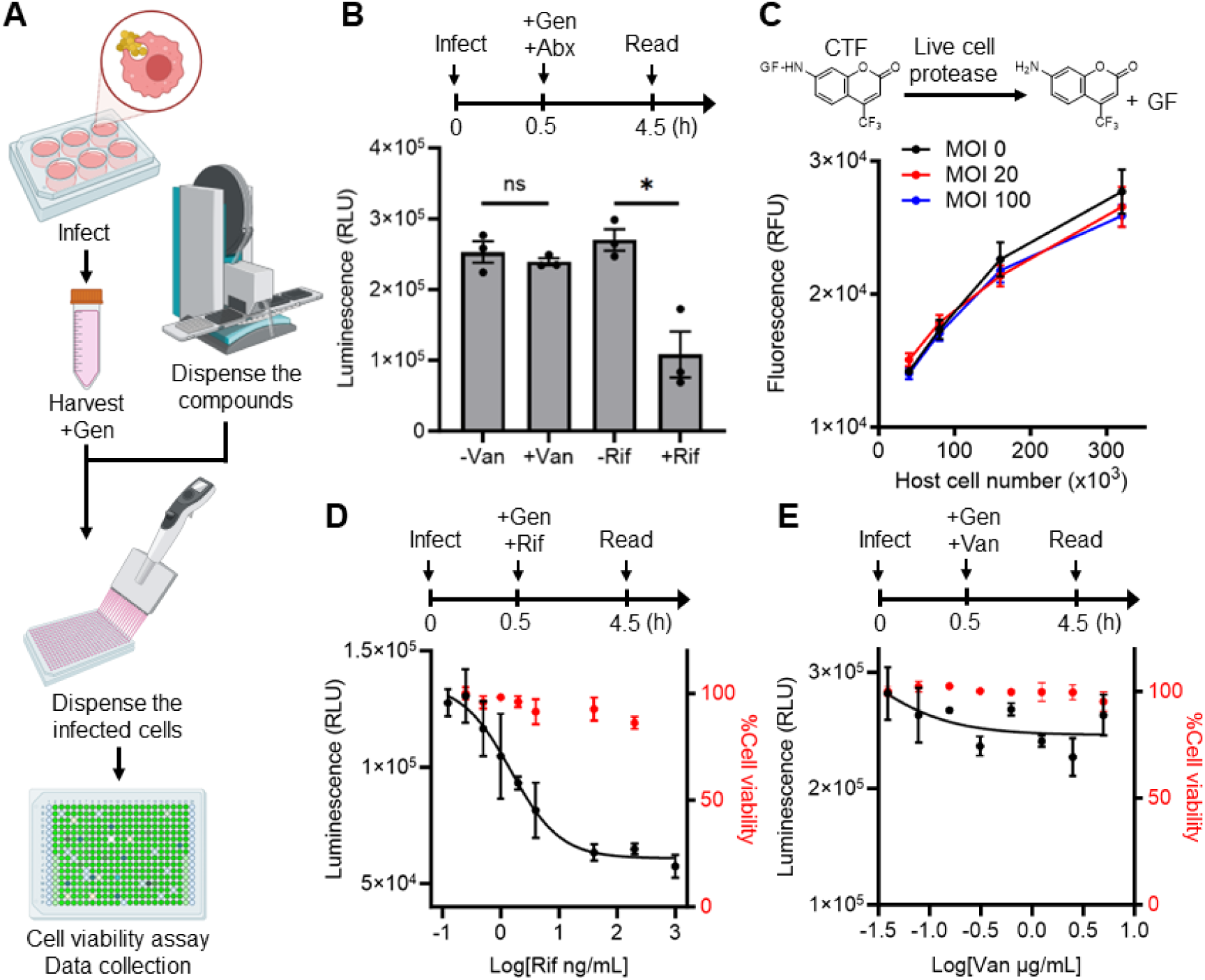
An established high-throughput screen allows for identifying compounds that modulate the energy state of intracellular *S. aureus*. (**A**) Schematic representation of the constructed compound screen for energy modulators of the intracellular bacteria. (**B**) The intracellular *S. aureus* activity can be probed using a bioluminescent MRSA strain JE2–lux. RAW 264.7 macrophages were infected with JE2– lux and treated with 20 µg/mL vancomycin (Van) or 10 µg/mL rifampicin (Rif) for 4 h before luminescence detection. Gentamicin (Gen; 50 µg/mL) was also added to eliminate the extracellular bacteria. Only Rif can penetrate the host plasma membrane and target the intracellular bacteria. Representative data of three biological replicates is shown (n = 3). *p<0.05; ns, not significant (unpaired t-test). The bars represent mean ± SEM. (**C**) The host cell viability can be monitored using CellTiter-Fluor (CTF). The enzymatic reaction reflected the number of viable RAW 264.7 cells (Pearson correlation coefficient r >0.97, p<0.05) and was not interfered with the presence of *S. aureus*. The multiplicities of infection (MOIs) of 0 (black), 20 (red) and 100 (blue) are shown (n = 3). (**D** and **E**) Dose response curves for Rif (**D**) and Van (**E**) inhibition of intracellular *S. aureus* activity (black circles). The host cell viability was simultaneously measured (red circles). Representative data of three biological replicates is shown (n = 3). The bars represent mean ± SEM. Assay schematics are shown above plots.

To identify energy modulators of intracellular *S. aureus* as antibiotic adjuvants, macrophages infected with JE2–lux were harvested from media containing gentamicin to eliminate extracellular bacteria, and then dispensed into 384-well plates with various compounds (**Figure 2A**). Automatic compound dispensing was performed using a high-throughput mosquito LV liquid dispenser. The bioluminescent signals and host cell viability were measured immediately following a 4-hour incubation. A total of >4,700 compounds that share structural similarity to kinase inhibitors and comply with the Lipinski’s rule of five were screened (Supplementary figure 1). From this kinase-targeted library, 77 compounds significantly reduced the bioluminescent signal to a level lower than that of the rifampicin-treated control (three standard deviations (SD) below the control). This signal reduction may indicate a decreased bacterial burden or a lower energy state of the intracellular bacteria. In this screen, we did not observe any changes in persister frequencies among 32 of the 77 compounds (42%) that exhibited the lowest signals.

Intriguingly, we identified 45 compounds that increased the bioluminescent signal by more than 1.5-fold, with no detectable cytotoxicity (**Figure 3A**). We found that the top candidate hit **KL1** (PubChem CID: 2881454) consistently increased the bioluminescent signal of intracellular *S. aureus* (Supplementary figure 2A). This elevated signal was not a result of gentamicin-killed extracellular bacteria, the host macrophages, or the compound itself (Supplementary figure 2). Importantly, co-administration of **KL1** sensitized intracellular MRSA to antibiotics that penetrate the mammalian plasma membrane, including rifampicin and moxifloxacin, by up to 10-fold (**Figure 3B**). Treatment with **KL1** alone did not induce any bacterial outgrowth, underscoring its ability to elevate the energy state of intracellular *S. aureus*. The other 44 compounds did not exhibit the same adjuvant activity at 10 µM, suggesting the minimum energy threshold for sensitization. To ensure its reliability, we repeated the experiments using **KL1** purchased from two independent suppliers and one synthesized in-house. In all cases, **KL1** increased intracellular *S. aureus* activity and enhanced the efficacy of rifampicin in killing the intracellular bacteria (Supplementary figure 3). This enhanced killing was observed across multiple laboratory strains and clinical isolates, and it was proportional to **KL1**’s concentration (**Figure 3C–D** and Supplementary figure 4). Furthermore, **KL1** retained its ability to synergize with rifampicin even when the intracellular bacteria were pretreated with the antibiotic, suggesting that low-energy persisters may be resuscitated and sensitized to antibiotic treatment (**Figure 3E**). Together, we demonstrated the feasibility of pharmaceutically sensitizing the intracellular *S. aureus* persisters to antibiotics and confirmed that screening these sensitizers using our newly established semi-automated high-throughput platform is viable.

**Figure 3.**
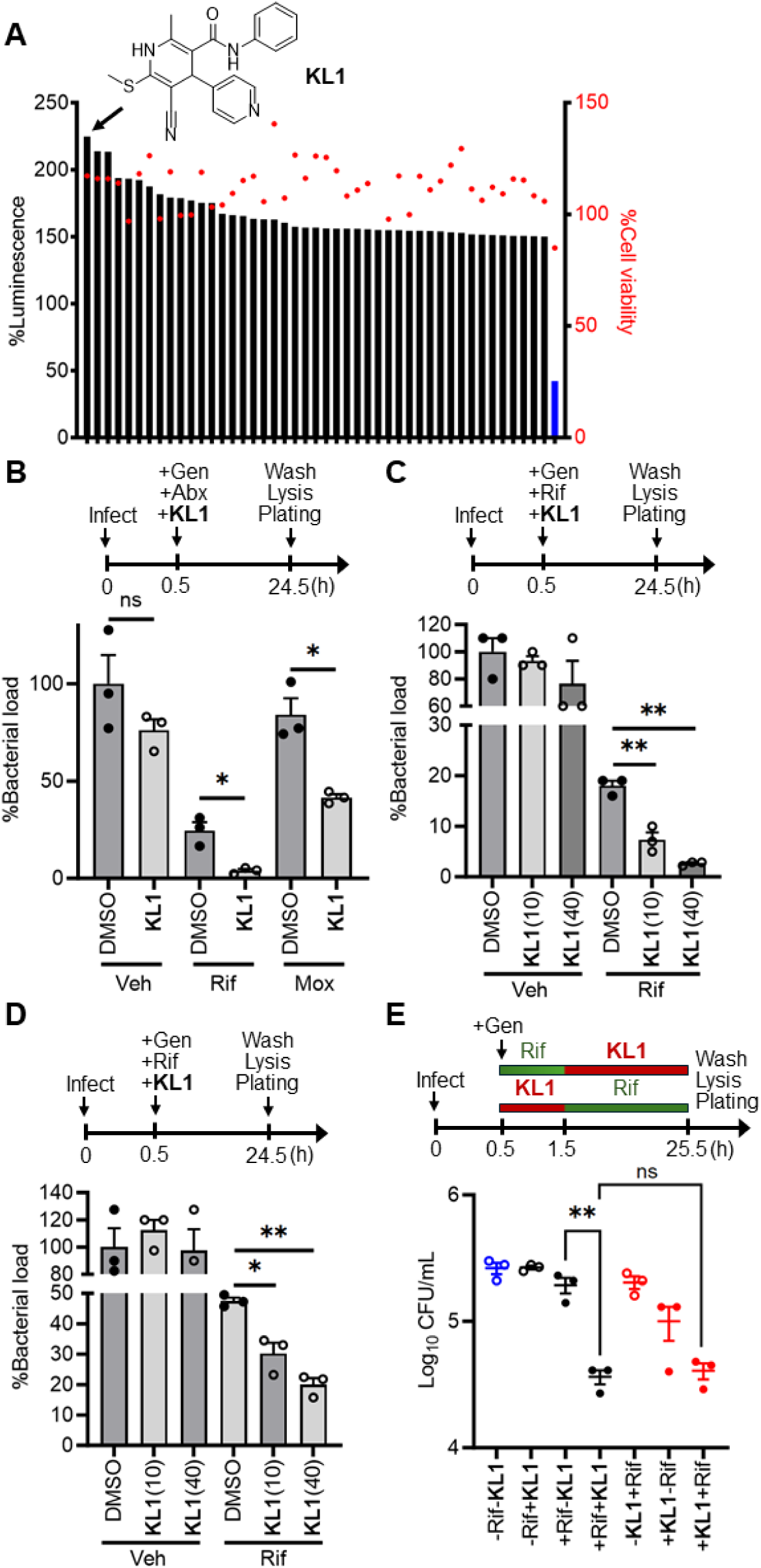
Screening a kinase-targeted compound library identifies a small molecule that sensitizes intracellular *S. aureus* to antibiotic killing. **(A)** Forty-five compounds (10 µM) exhibited >1.5-fold increase in the luminescence signal (black bars) comparing to the vehicle control (0.1% DMSO). No cytotoxicity was observed in the 4-h treatment window (red circles). The chemical structure of the top candidate compound **KL1** is shown. The rifampicin-treated group was included as a reference (blue bar). (**B**) **KL1** (open circles) facilitates rifampicin (Rif; 10 µg/mL) and moxifloxacin (Mox; 50 µg/mL) killing of the intracellular MRSA. Gentamicin (Gen; 50 µg/mL) was added to eradicate the extracellular bacteria. The numbers of survivor bacteria were normalized to the untreated control (no antibiotics (Abx), no **KL1**, Gen-only) (n = 3). (**C** and **D**) The **KL1**-mediated synergism was detected in different clinical isolates with the high persister (**C**) and low persister (**D**) phenotypes characterized in Figure 1A. A dose-dependent sensitization was observed with 0–40 µM **KL1** in these isolates (n = 3). (**E**) Pre-exposure to Abx does not affect the sensitization by **KL1**. Infected RAW 264.7 cells were treated with either 10 µg/mL Rif (black circles) or 40 µM **KL1** (red circles) for 1 h, followed by the addition of **KL1** and Rif, respectively. The untreated group (blue circles) was included as a negative control (n = 3). Assay schematic above plot indicates the orders of the treatment regimens. *p<0.05; **p<0.01; ns, not significant (unpaired t-test). The bars represent mean ± SEM.

### Compound KL1 exhibits adjuvant activity in a mouse model for *S. aureus* sepsis and a cell-based *Salmonella* infection model

Next, we evaluated the in vivo efficacy of **KL1** using our *S. aureus* bacteremia model (15). Briefly, C57BL/6J mice were intravenously injected with *S. aureus* to induce a systemic infection. At 6 hours post-infection (hpi), mice were administered 10 mg/kg rifampicin once daily (q.d.), either alone or in combination with 100 mg/kg **KL1** twice daily (b.i.d.), via intraperitoneal injections (**Figure 4A**). After a 48-hour treatment regimen, the livers and spleens were harvested, and the tissue homogenates were plated to determine the number of bacteria that survived the antibiotic treatment. Our data showed that combined administration of **KL1** and rifampicin further reduced the bacterial burden in both the liver and spleen, compared to rifampicin alone (**Figure 4B** and **C**). This animal experiment was replicated on separate days using both male and female mice from different litters, which each group consisting of 6–8 mice, to ensure reproducibility.

**Figure 4.**
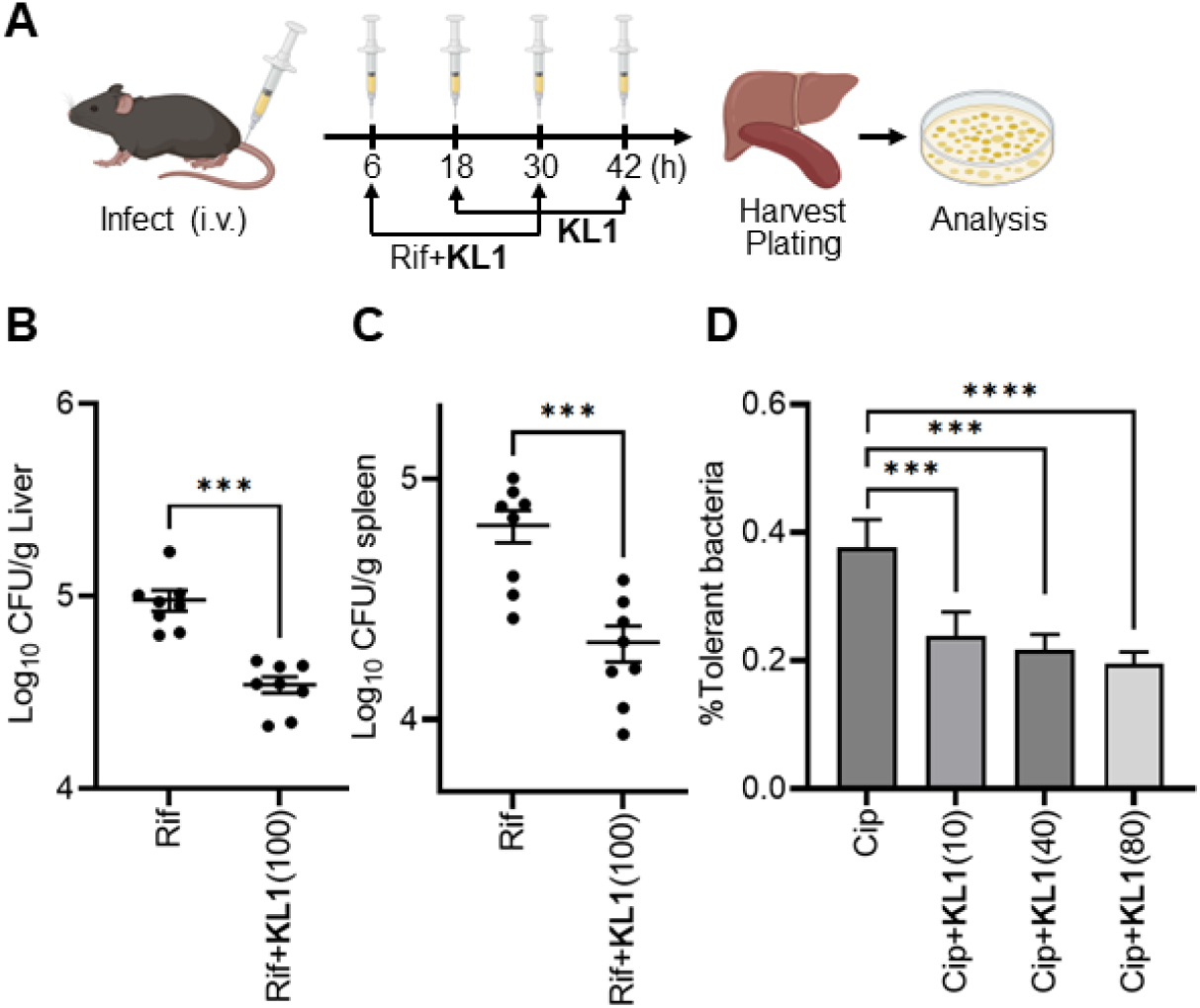
The identified compound KL1 enhances antibiotic killing of *S. aureus* in a murine bacteremia model and *Salmonella Typhimurium* in macrophages. (**A**) Schematic representation of the murine model. C57BL/6J mice were infected via intravenous (i.v.) tail vein injections and intraperitoneally treated with 10 mg/kg rifampicin (Rif) (q.d.) with and without 100 mg/kg **KL1** (b.i.d.) at 6 hours post-infection (hpi) for 2 days. The organs were harvested and homogenized to enumerate the numbers of survivor bacteria (CFU). Co-administration of **KL1** and Rif reduced the bacterial burden in the liver (**B**) and spleen (**C**). Representative data of two independent experiments is shown (n = 8). A total of 14 mice per treatment group were examined. ***p<0.001 (unpaired t-test). (**D**) **KL1** facilitates ciprofloxacin (Cip; 5 µg/mL) killing of intracellular *Salmonella Typhimurium*. The numbers of survivor bacteria were normalized to the input CFUs (T_0_) (n = 7–8). ***p<0.001; ****p<0.0001 (two-way ANOVA). The bars represent mean ± SEM.

Since **KL1** may or may not directly target the bacteria and may instead modulate host– pathogen interactions, we explored the possibility that **KL1** might also affect other intracellular bacteria. Here, we opted to investigate the gram-negative bacterium *Salmonella typhimurium* because it also survives within macrophages and forms antibiotic-tolerant persister cells (18). Strikingly, **KL1** exhibited adjuvant activity when combined with ciprofloxacin and enhanced the killing of *Salmonella typhimurium* in macrophages (**Figure 4D**). This data suggests that **KL1** may serve as a potential broad-acting antibiotic adjuvant against intracellular bacteria.

### KL1 does not directly act on *S. aureus*

Since the **KL1**-mediated synergy was likely energy-dependent, we utilized a mutant MRSA strain lacking all four glucose transporters (ΔG4) to test if the bacterial central metabolism was crucial for **KL1**’s activity (35). We found that the adjuvant activity of **KL1** against intracellular *S. aureus* was substantially attenuated in the absence of functional glucose uptake (Supplementary figure 5). This finding aligned with the hypothesis that **KL1** enhanced antibiotic efficacy by modulating the energy state of intracellular bacteria and suggested that glucose utilization was essential for sensitization of the tolerant cells.

To deconvolute **KL1**’s mechanism of action, we interrogated whether **KL1** impacted extracellular *S. aureus* in planktonic cultures. We began by testing **KL1**’s activity in actively growing *S. aureus* at the mid-exponential phase (high energy state). In this case, no increase in energy was detected with 10 µM **KL1**, despite observing a moderate increase at the higher concentration of 40 µM (Supplementary figure 6A). Treatment with **KL1** at the higher concentration did not alter bacterial growth, nor did it affect persister frequency following rifampicin treatment (Supplementary figure 6B). Similarly, **KL1** did not affect the energy state or growth pattern of non-growing, stationary-phase bacteria (low energy state) in the presence or absence of antibiotics (Supplementary figure 6C and D).

We subsequently employed Seahorse analysis to investigate whether **KL1** altered the metabolic activity of extracellular *S. aureus*. This analysis measures real-time changes in cellular metabolism and has been applied to investigate bacterial respiration in *S. aureus* (36). Our data clearly showed that **KL1** had no effect on the oxygen consumption rate (OCR) and extracellular acidification rate (ECAR) of *S. aureus* in planktonic cultures (Supplementary figure 7A and B). Additionally, we directly measured the relative ATP levels in the bacteria and did not observe any changes across multiple *S. aureus* strains (Supplementary figure 7C–E). Collectively, our data strongly suggested that **KL1** did not directly target the bacteria but may perturb the host–pathogen interaction by altering the intracellular environment.

### KL1 modulates the host cells and reduces the level of reactive species

To elucidate **KL1**’s mode of action, we profiled the transcriptomic changes of *S. aureus*-infected macrophages in response to the treatment using bulk RNA-sequencing. In comparison with the vehicle control, **KL1** treatment resulted in the up-regulation of 24 genes and the down-regulation of 90 genes (**Figure 5**). The **KL1**-mediated changes were enriched in inflammatory responses, cytokine production and macrophage activation. However, the treatment did not appear to obliterate the host immune response, and treatment with **KL1** alone did not cause bacterial outgrowth. Intriguingly, multiple host genes involved in regulating reactive oxygen and nitrogen species (ROS/RNS) were identified (**Figure 5B and C**). Important genes promoting ROS/RNS production, such as *Thbs1*, *S100a8*, *Nos2* and *Maf*, were significantly down-regulated, while a glutathione S-transferase gene *Gsta2*, involved in the cellular anti-oxidative defense was up-regulated (37, 38). As such, **KL1** may alleviate the oxidative and nitrosative stress in the intracellular niche, which aligns with previous findings that ROS/RNS can induce antibiotic tolerance and persister cells in intracellular *S. aureus* and various bacterial species (15, 16, 18–21, 39).

**Figure 5.**
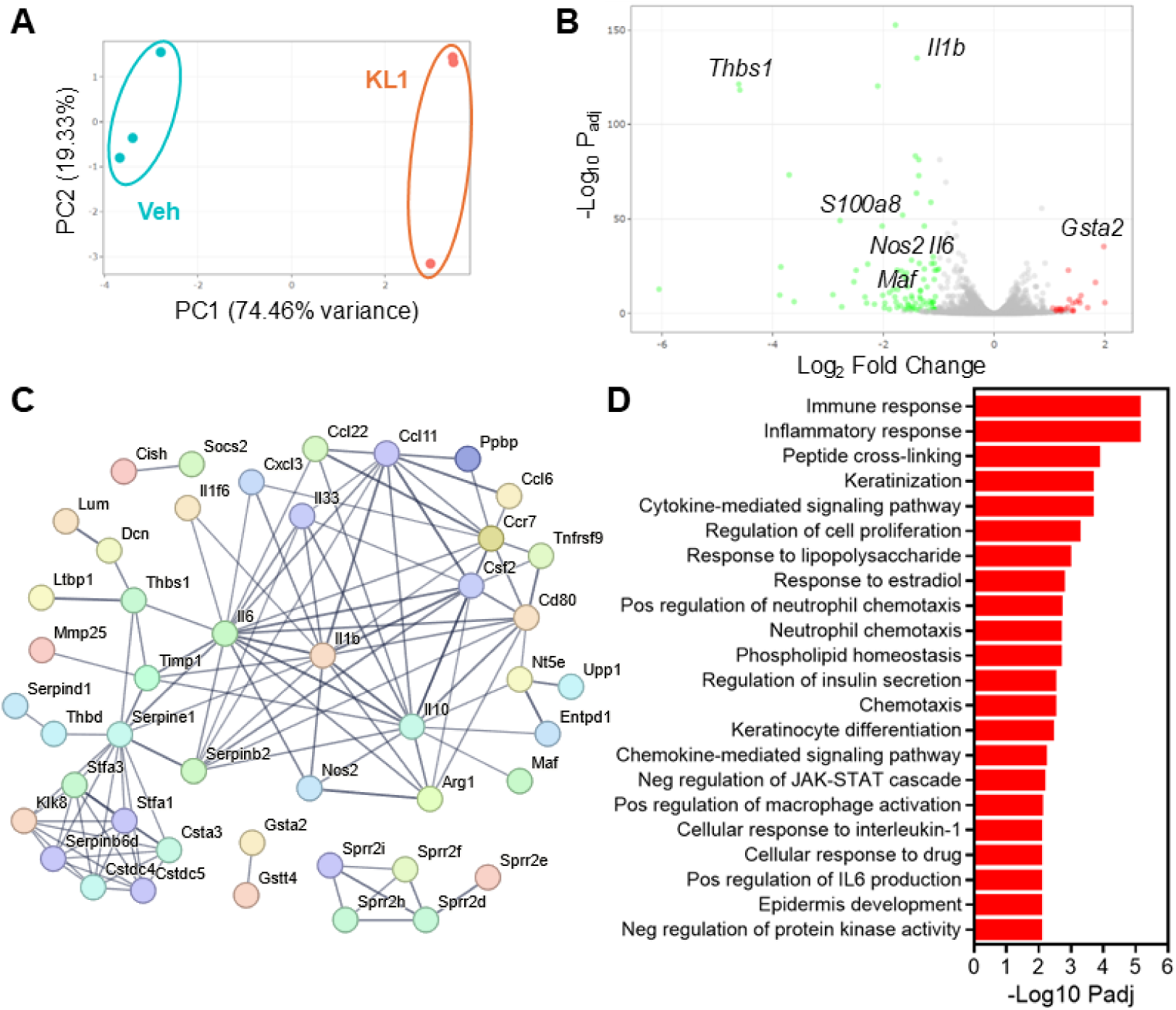
KL1 modulates the expression of host immune response genes regulating reactive oxygen and nitrogen species. Principle component analysis (**A**), differential expression (**B**) and String network (**C**) analysis illustrate the **KL1**-mediated transcriptional modulation in *S. aureus*-infected macrophages (n = 3). Genes associated with reactive species production are indicated. Edge thickness represents the confidence in the interactions. Interactions with high confidence (interaction score >0.7) are shown. (**D**) Gene ontology enrichment with an adjusted p-value <0.01 is shown.

Although the exact mechanism of **KL1** has not been determined, this compound has been included in hundreds of biological screens. Among these screens, **KL1** was found to be active in five bioassays targeting four different proteins (Supplementary table 3). EHMT2/G9a, in particular, stood out as a potential candidate target of **KL1** given that this epigenetic regulator is an intracellular protein expressed in immune cells, including macrophages. Additionally, the EC_50_ values of **KL1** (25 µM) determined in the previous EHMT2/G9a inhibition assay (PubChem AID: 504332) aligned with the adjuvant activity of **KL1** (10–40 µM) identified in this study. Intriguingly, EHMT2/G9a plays a key role in the intracellular innate immunity and modulates macrophage polarization through the regulation of inflammatory genes, such as *Il6* and *Il1b* (40–44). It can also regulate NF-κB and JAK/STAT signaling (43). This downstream regulation is partly reminiscent of our findings in **KL1**-mediated gene expression shift (**Figure 5**). To interrogate EHMT2/G9a as a potential target, we first utilized the highly selective EHMT inhibitor BIX-01294 to assess whether EHMT2/G9a influences antibiotic susceptibility of intracellular *S. aureus*. Surprisingly, we found that inhibition of EHMT2/G9a also elevated the energy state of intracellular *S. aureus* and synergized with rifampicin in a dose-dependent manner, similar to the effect observed with the identified compound **KL1** (**Figure 6A** and **B**). Given the role of EHMT2/G9a in modulating inflammatory responses and our findings that **KL1** alters the expression of genes involved in ROS/RNS production, we hypothesized that **KL1** targets EHMT2/G9a and alleviates the production of ROS/RNS, thereby facilitating the antibiotic killing of intracellular bacteria. To test this hypothesis, we employed two probing methods with distinct detection mechanisms to measure reactive species. Specifically, a chemiluminescent probe L-012 and a fluorescent probe fluorescein-boronate (Fl-B) were applied (16, 45). Using both methods, we consistently observed that **KL1** at 40 and 100 µM significantly reduced the levels of ROS/RNS in *S. aureus*-infected macrophages at 4- and 8-hours post-treatment (**Figure 6C–D** and Supplementary figure 8A and C). Notably, treatment with the EHMT2/G9a inhibitor BIX-01294 and the antioxidant control butylated hydroxyanisole (BHA) both resulted in a similar decrease in ROS/RNS. Furthermore, **KL1** was able to lessen the levels of ROS/RNS in uninfected macrophages, highlighting its ability to adjust the host environment (Supplementary figure 8B and D).

**Figure 6.**
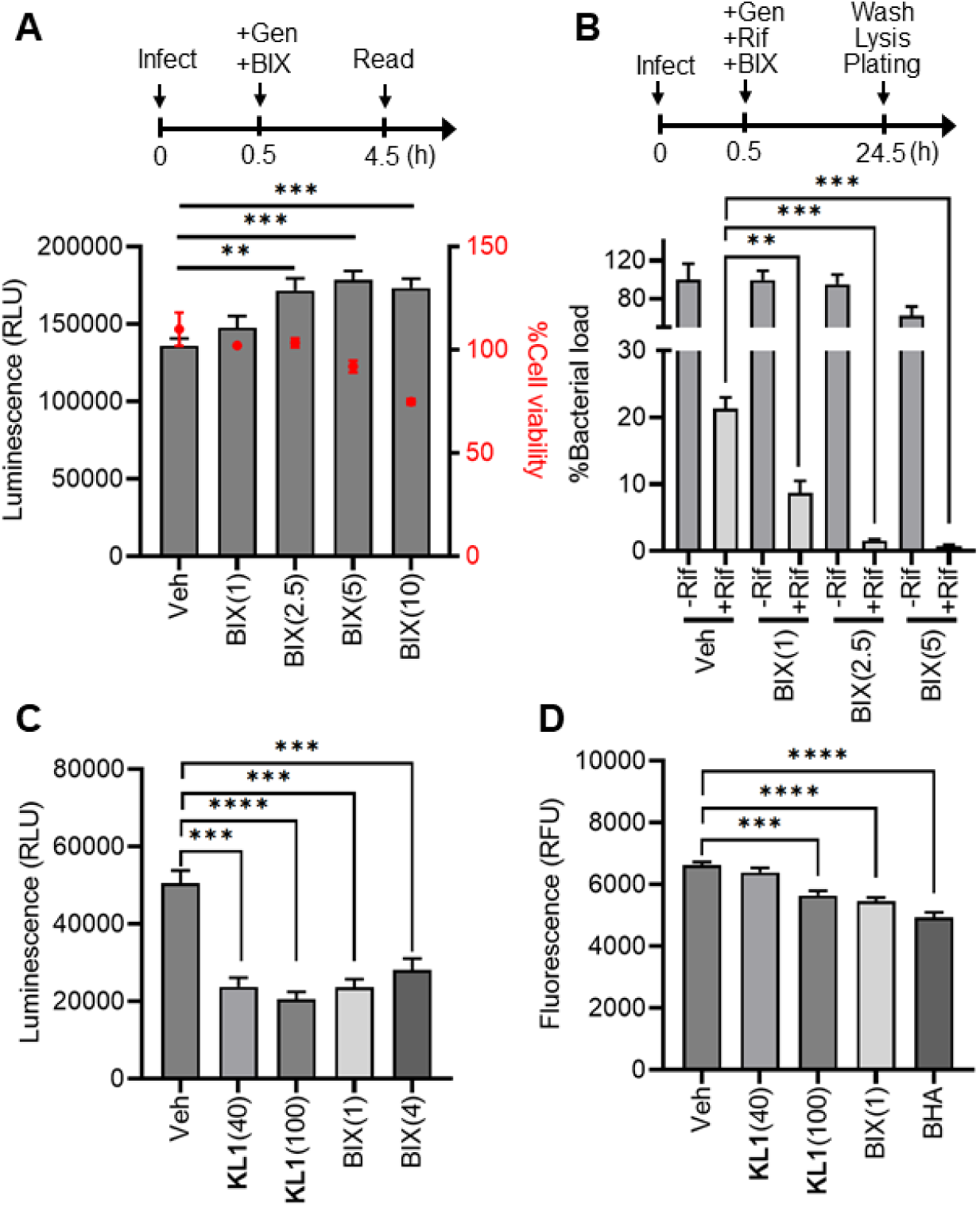
KL1 reduces the production of reactive species upon infection. (**A**) Inhibition of G9a/EHMT2 phenocopies the **KL1**-mediated sensitization in intracellular *S. aureus*. A selective G9a inhibitor, BIX-01294 (BIX; 2.5–10 µM), increased the bioluminescence signal (grey bars) of strain JE2–lux comparing to the vehicle control (Veh, 0.1% DMSO). The relative cytotoxicity of BIX was examined (red circles). Representative data of three independent experiments is shown (n = 5). (**B**) The G9a inhibitor increases rifampicin (Rif; 10 µg/mL) killing of the intracellular MRSA in a dose-dependent manner (1–5 µM). Gentamicin (Gen; 50 µg/mL) was added to eradicate the extracellular bacteria. The numbers of survivor bacteria were normalized to the untreated control (no Rif, no BIX, Gen-only) (n = 3). Both **KL1** (40 and 100 µM) and BIX (1 and 4 µM) reduced ROS production at 4 hours post-infection (hpi). A chemiluminescent probe L-012 (**C**) and a fluorescent probe fluorescein-boronate (Fl-B) (**D**) were utilized to quantify reactive species in infected macrophages. An antioxidant butylated hydroxyanisole (BHA; 20 µM) was included as a control. Representative data of three independent experiments are shown (n = 5). Assay schematics are shown above plots. **p<0.01; ***p<0.001; ****p<0.0001 (unpaired t-test). The bars represent mean ± SEM.

### Inactive analog of KL1 losses the ability to modulate the level of reactive species

To provide further insight into the medicinal chemistry, we conducted a brief structure– activity relationship study using six analogs of **KL1** and tested how different chemical modifications interfered with their biological activity (**Figure 7A**). The substitution of carboxamide at the alkyl terminal (KL2, KL4 and KL5) or removal of the nitrile group from the dihydropyridine ring (KL6) led to moderate potency loss in sensitizing intracellular bacteria to rifampicin (**Figure 7B**). The hexyl substitution on sulfur (KL3) resulted in a high cytotoxicity against host macrophages, rendering a poor therapeutic index for targeting intracellular bacteria. Notably, the 1,4-dihydropyridine ring oxidized analog (KL7) completely lost the adjuvant activity, presumably resulted from the altered electronic structure, decreased conformational flexibility or loss of hydrogen bonding capacity (**Figure 7B**). Importantly, this inactive analog KL7 also lost the ability to reduce host-derived reactive species in both infected and uninfected macrophages (**Figure 7C** and Supplementary figure 9). Altogether, these results strongly support the hypothesis that **KL1** enhances antibiotic killing of intracellular bacteria by reducing oxidative and/or nitrosative stress in the intracellular milieu, presumably by targeting the epigenetic regulator EHMT2/G9a (**Figure 8**).

**Figure 7.**
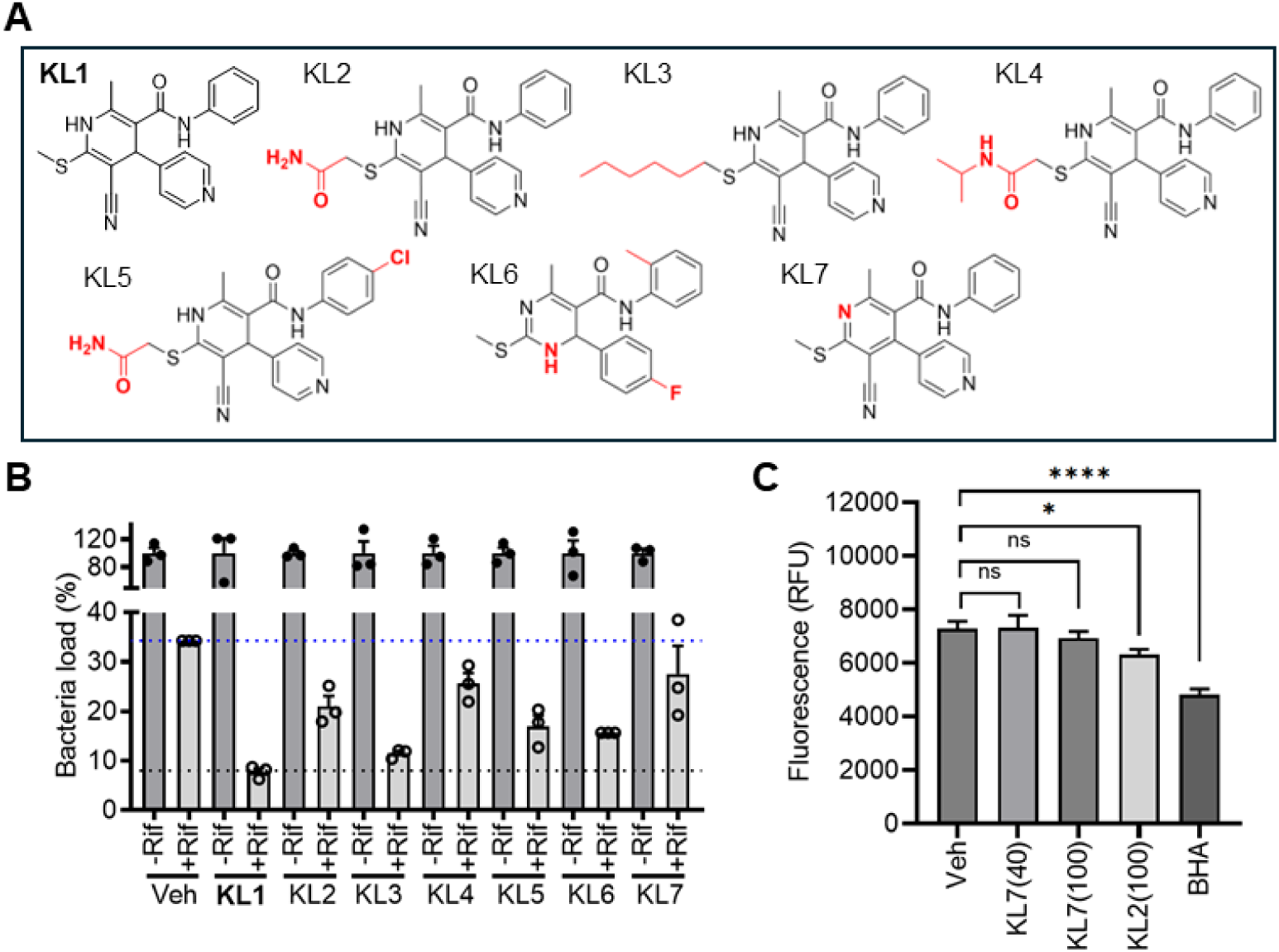
KL1 analogs with attenuated adjuvant activity loss the ability to regulate reactive species production. (**A**) Chemical structures of **KL1** and its analogs. Modified functional groups were indicated in red. (**B**) **KL1** analogs (40 µM) exhibited differential activities in sensitizing intracellular *S. aureus* to antibiotic killing. Infected cells were treated with (open circle) and without (solid circle) rifampicin (Rif; 10 µg/mL) and various analogs. Gentamicin (50 µg/mL) was added to eliminate the extracellular bacteria. The relative bacterial loads were normalized to the corresponding controls (-Rif). (**C**) The inactive analog KL7 (40 and 100 µM) and KL2 (100 µM) with reduced adjuvant activity showed no and minimum effect on ROS/RNS production in infected macrophages, respectively. DMSO (Veh) and an antioxidant butylated hydroxyanisole (BHA; 20 µM) were included as controls. Representative data of three biological replicates is shown (n = 3). *p<0.05; ****p<0.0001; ns, not significant (unpaired t-test). The bars represent mean ± SEM.

**Figure 8.**
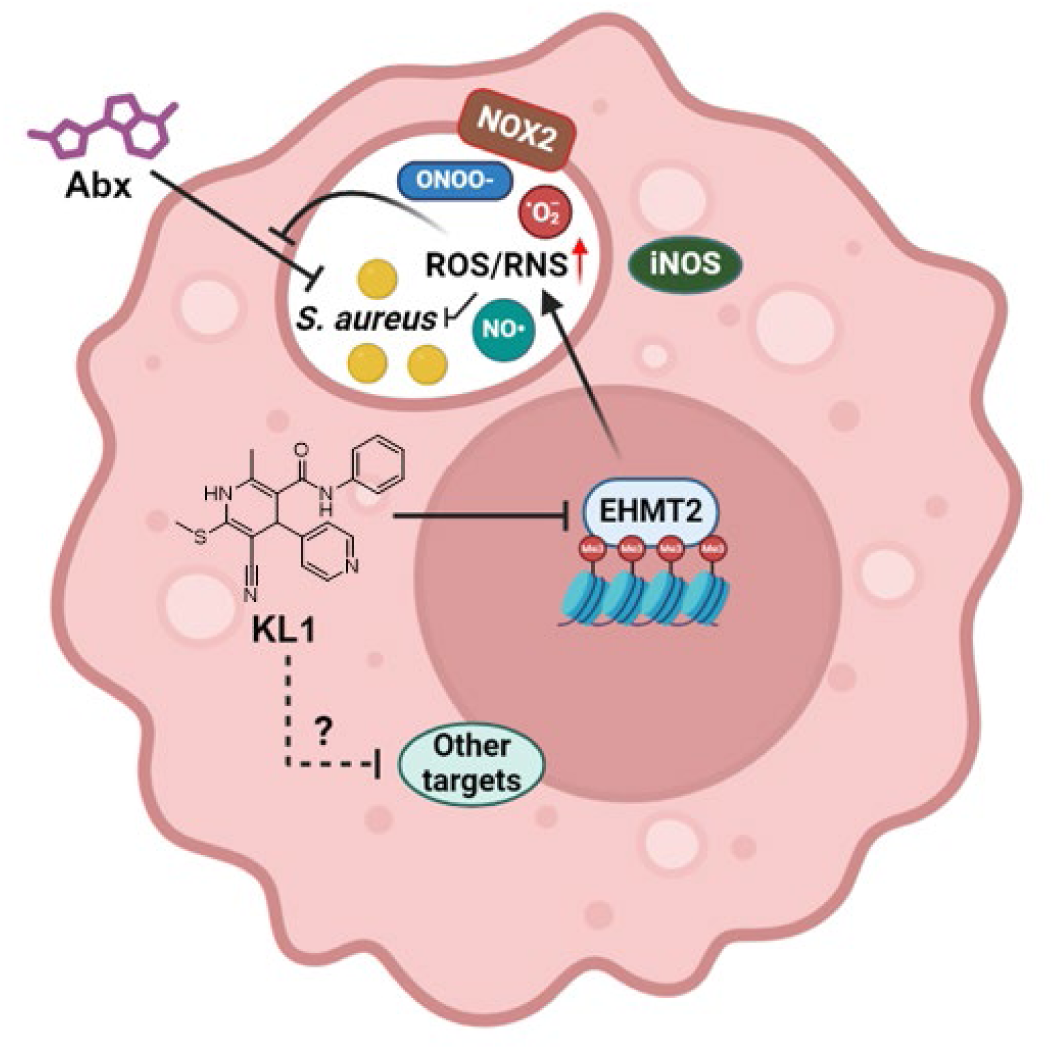
Mechanistic model of KL1-mediated sensitization of intracellular *S. aureus*. Host-derived reactive oxygen and nitrogen species (ROS/RNS) help contain intracellular *S. aureus* infections but also antagonize antibiotics by collapsing the bacterial metabolism and forcing the bacteria into antibiotic-tolerant persister cells. The identified compound **KL1** sensitizes intracellular bacteria to antibiotic killing by reducing the production of reactive species upon infection, presumably by targeting the G9a/EHMT2 activity.

## Discussion

The ability of bacterial populations or sub-populations of persister cells to survive high concentrations of bactericidal antibiotics for extended periods, despite lacking resistance mutations, is likely a major factor contributing to treatment failure. Approximately 20–30% of patients with *S. aureus* bacteremia fail to clear the infection despite receiving appropriate antibiotic therapy (46–48). The weak correlation between in vitro antimicrobial susceptibility and clinical outcomes highlights the importance of considering the relevant physiological context when addressing the poor antibiotic efficacy in clinical settings (6).

Devising novel strategies to target antibiotic tolerant persister cells has drawn increasing attention over the years (14, 49–51). Conceptually, the eradication of persister cells can be achieved either by directly targeting the persister cells or by reverting them to a metabolically active state, thereby making them vulnerable to antibiotics. The intracellular environment is a well-established reservoir for antibiotic tolerant persister cells (15–19). In this study, we demonstrated the potential of altering the host microenvironment to sensitize intracellular persisters to antibiotics and improve treatment outcomes. Although different stress stimuli can lead to the formation of antibiotic tolerant persister cells, most produce a metabolically indolent state that can withstand antibiotic challenge. We therefore employed an ATP-dependent bioluminescent reporter to probe the energy levels of intracellular bacteria as part of a phenotypic screen. When combined with a host cell viability measurement and CFU determination, this platform allowed us to identify molecules that activate intracellular persister cells without compromising host viability or inducing expansion of the pathogen. Given that the low-energy, low-metabolic activity state is central to antibiotic tolerance and persistence in multiple bacterial species, this screening may be adapted for other intracellular pathogens across various host cell types and will be agnostic to the mechanism responsible for the induction of antibiotic tolerance (52–55).

Intriguingly, our lead compound **KL1** significantly reduced ROS/RNS production in macrophages, sensitized intracellular *S. aureus* and *Salmonella* persister cells to antibiotic treatment, and enhanced antibiotic efficacy in a murine infection model. Transcriptomic analysis further support a mechanism associated with an altered inflammatory response and reduced ROS/RNS production. These observations are in line with a previous biochemical activity screen in which **KL1** inhibited the activity of EHMT2/G9a that epigenetically regulates a wide range of biological processes, including macrophage polarization (40–44). Consistently, inhibition of EHMT2/G9a mirrored the activity of **KL1** and caused sensitization of intracellular *S. aureus*, implicating that EHMT2/G9a may be the primary target of **KL1** (**Figure 8**).

Together, these findings strongly support that: 1) Targeting host pathways can render intracellular persisters more susceptible to antibiotics; 2) ROS/RNS are key drivers of antibiotic tolerance in vivo; 3) Inhibiting ROS/RNS production can sensitize persister cells across diverse bacterial species; and 4) Sensitizing intracellular persisters leads to improved antibiotic efficacy against active infections. This work underscores the potential of using host-directed therapeutics, in conjunction with antibiotics, to enhance treatment outcomes. Continued development of our screening platform promises identification of additional host-targeted drugs that sensitize intracellular persisters to antibiotics. Ongoing studies on **KL1**’s mechanism of action, along with medicinal chemistry optimization and in vivo testing in combination with various antibiotics against multiple pathogens, including *M. tuberculosis*, will be critical in determining the full therapeutic potential of this host-directed adjuvant in eradicating deep-seated infections.

## Materials and methods

### General materials

Antibiotics including rifampicin (Rif), moxifloxacin hydrochloride (Mox), gentamicin sulfate (Gen) were purchased from Fisher Scientific with >95% purity. Vancomycin hydrochloride (Van) (>99% purity) was purchased from Alfa Aesar. Compound **KL1** was obtained from multiple sources (ChemBridge Corporation, Enamine and synthesized in-house (Supplementary Information)) with >90% purity to validate the fidelity of this compound. Analogs KL2–6 were purchased from ChemDiv, ChemBridge Corporation and Vitas M Chemical Limited (>90% purity). Analog KL7 was synthesized in-house with >90% purity (Supplementary Information). The G9a inhibitor, BIX-01294 (>98% purity), was purchased from MedChemExpress. RAW 264.7 macrophages were obtained from ATCC (TIB-71) and distributed by the Tissue Culture Facility at the University of North Carolina at Chapel Hill. RAW 264.7 cells were maintained in complete DMEM (Dulbecco’s Modified Eagle Medium (Gibco) supplemented with 10% (vol/vol) heat-inactivated fetal bovine serum (FBS) (Avantor Seradigm, premium grade), 2 mM L-glutamine (Gibco), 1x non-essential amino acids solution (Gibco) and 1 mM sodium pyruvate (Gibco)) below 18 passages without reaching more than 90% confluency. Bone marrow-derived macrophages (BMDMs) were isolated from C57BL/6J mice (The Jackson Laboratory; strain number 000664) and cultivated in complete DMEM as previously described (56). Immortalized BMDMs were generated using a CRE-J2 retroviral infection method (57).

Briefly, C57BL/6J mouse-derived BMDMs were cultured in 50% (vol/vol) L929 fibroblast-conditioned medium and infected with CRE-J2 retrovirus on day 5 and day 7. Transduced cells were continuously cultured in the conditioned medium, with the concentration gradually reduced to 20% conditioned medium. The immortalized BMDMs were maintained in RPMI 1640 (Corning) supplemented with 2 mM L-glutamine, 1x Penicillin-Streptomycin (GenClone), 10% (vol/vol) heat-inactivated FBS (R&D Systems) and 20% conditioned medium.

### Bacterial strains and growth conditions

*S. aureus* strains LAC (USA300), HG003, JE2 (USA300), JE2–lux (JE2*lux*ABDCE) (kindly provided by Roger Plaut, Center for Biologics Evaluation and Research, Silver Spring, United States) (34), ΔG4 (kindly provided by Anthony Richardson, University of Pittsburgh, Pittsburgh, United States) (35) and the *S. aureus* bacteremia isolates (16) were routinely cultured in tryptic soy broth (TSB) (Fisher Scientific) at 37 °C and 225 rpm unless otherwise stated. The clinical isolates were obtained under an IRB exemption from a preexisting collection. *Salmonella enterica serovar Typhimurium* strain 14028S was cultured in LB broth (Lennox) at 37 °C with shaking.

### Minimum inhibitory concentration assay

Stationary-phase bacterial cultures were diluted at 1:1,000 in Mueller–Hinton broth (MHB) (Oxoid) containing serially diluted Rif (0–0.2 µg/mL), Mox (0–20 µg/mL) or Van (0–16 µg/mL) in 96-well assay plates (Corning) in triplicates (200 µL/well). Plates were sealed with Breathe-EASIER membranes (Diversified Biotech) and incubated statically at 37 °C for 24 h. The minimum inhibitory concentrations (MICs) were determined by the absence of bacterial growth. Three independent assays were performed to ensure the reproducibility.

### Construction of the inducible reporter strains

The plasmid pALC2084 containing a tetracycline (Tet)-inducible GFP-expressing cassette (58) (kindly provided by Ambrose Cheung, Dartmouth College, Hanover, United States) was transformed into a *S. aureus* intermediate strain RN4220 using a Gene Pulser Xcell electroporation system (Bio-Rad) as previously described (59). The plasmid was subsequently purified and transformed into the recipient *S. aureus* strain HG003. Single colonies were isolated and cultured in TSB containing 10 µg/mL Cam and 0–2 µM anhydrotetracycline hydrochloride (aTc) (Sigma) for 2–3 h. The induction of GFP expression was verified at an excitation and emission wavelength of 475 nm and 508 nm using a Synergy H1 microplate reader (BioTek). To construct the mKate reporter plasmid, the DNA fragment of mKate2 was amplified from pRN10 (Addgene plasmid: 84454) (60) using a specific primer pair (Supplementary table 1) and Q5 High-Fidelity DNA Polymerase (New England Biolabs). The PCR product was purified using a MinElute Reaction Cleanup kit (Qiagen). A Tet-inducible expression vector pRMC2 (Addgene plasmid: 68940) (61) and the mKate DNA fragments were subjected to KpnI-HF and EcoRI-HF (New England Biolabs) enzymatic digestion and agarose gel electrophoresis. The DNA fragments were gel purified using a QIAquick PCR purification kit (Qiagen) and underwent ligation with T4 DNA ligase (New England Biolabs) at 16 °C for 16 h. The ligation product was transformed into MAX Efficiency DH5α competent cells (ThermoFisher Scientific) according to the manufacturer’s instruction. Site-directed mutagenesis was performed using a QuikChange kit (Agilent Technologies) and a primer pair (Supplementary table 1) to incorporate an upstream ribosomal binding site. The final pRMC2-mKate construct was verified by restriction digest mapping and Sanger sequencing. The plasmid pRMC2-mKate was transformed into strain RN4220, followed by the recipient strain HG003 as described above. The aTc-induced mKate expression was confirmed by measuring at 633 nm with excitation at 588 nm using a Synergy H1 microplate reader.

### Antibiotic survival assays

For *S. aureus* planktonic cultures, 16- to 18-h stationary cultures were diluted at 1:100 in MHB and incubated at 37 °C and 225 rpm for 3 h to reach the mid-exponential phase. The bacterial cultures were then treated with 10 µg/mL Rif and/or 0–40 µM **KL1** at 37 °C and 225 rpm for 24 h. The bacteria were washed in one volume of sterile 1% NaCl three times at 20,000 *g* for 5 min, followed by resuspending in one volume of 1% NaCl. The bacterial suspensions were 10-fold serially diluted and plated on tryptic soy agar (TSA) to enumerate the number of survivor *S. aureus*. The frequencies of tolerant bacteria were calculated by normalizing the numbers of survivor bacteria to the numbers of input bacteria (T_0_).

To assess the intracellular *S. aureus* persister frequencies, RAW 264.7 macrophages or BMDMs were seeded in Costar 24-well tissue culture-treated plates (Corning) (500 µL of 4×10^5^ cells per well) in complete MEM (Minimum Essential Medium (Gibco) supplemented with 10% (vol/vol) heat-inactivated fetal bovine serum and 2 mM L-glutamine) at 37 °C and 5% CO_2_ for 16 h. Macrophages were infected with *S. aureus* strains at a multiplicity of infection (MOI) of 10 by centrifugation at 1,000 *g* for 2 min, followed by incubation at 37 °C for 35 min. After removal of the spent medium, infected cells were incubated in 500 µL complete MEM containing antibiotics (10 µg/mL Rif, 50 µg/mL Mox or 20 µg/mL Van) in the presence or absence of 0–40 µM **KL1** and its analogs (KL2–7) or 0–10 µM BIX-01294 in the presence of 50 µg/mL Gen at 37 °C and 5% CO_2_ for 24 h. The cells were washed three times in 1 mL PBS and lysed with 200 µL 0.5% (vol/vol) Triton X-100 (Fisher Scientific) at 37 °C for 5 min, followed by the addition of 800 µL PBS. The released intracellular *S. aureus* were 10-fold serially diluted and plated on TSA for enumeration. For the persister frequencies, the numbers of survivor bacteria were normalized to the corresponding numbers of intracellular bacteria before the antibiotic treatment (T_0_). For comparison between different treatment groups, the intracellular *S. aureus* loads were normalized against the untreated controls at the endpoint (24-h post-treatment).

For *Salmonella* infections, immortalized BMDMs were seeded in 12-well tissue culture-treated plates (4×10^5^ cells per well) in RPMI 1640 supplemented with 2 mM L-glutamine and 10% (vol/vol) heat-inactivated FBS at 37 °C and 5% CO_2_ for 24 h. The stationary-phase bacteria were opsonized with 8% (vol/vol) mouse serum in the RPMI 1640 medium for 20 min and applied to the immortalized BMDMs at an MOI of 15 as previously described (32). The plates were centrifuged at 110 *g* for 5 min and incubated at 37 °C and 5% CO_2_ for 30 min. Cells were washed once with PBS and incubated in RPMI 1640 medium containing 5 µg/mL ciprofloxacin hydrochloride (MP Biomedicals) in the presence of 10–80 µM **KL1** or 0.2% DMSO at 37 °C and 5% CO_2_ for 24 h. The cells were washed three times in 500 µL PBS and lysed with 0.2% (vol/vol) Triton X-100 at 37 °C for 10 min. The extracted intracellular *Salmonella Typhimurium* were plated on LB agar to determine the number of survivor bacteria. The persister frequencies were calculated by normalizing the final CFUs (T_24_) to the input CFUs (T_0_).

### Live cell microscopy

RAW 264.7 macrophages were seeded in a Nunc 8-well Lab-Tek chambered coverglass (ThermoFisher Scientific) (500 µL of 2–3×10^5^ cells per well) in complete MEM at 37 °C and 5% CO_2_ for 16 h. Cells were infected with the Tet-inducible GFP- or mKate-expressing *S. aureus* strains at an MOI of 20 (without Rif) or 100 (with Rif) by centrifugation at 1,000 *g* for 2 min and incubating at 37 °C for 35 min. Cells were washed once in 500 µL complete MEM and treated with and without 10 µg/mL Rif in the presence of 50 µg/mL Gen at 37 °C for 1.5 h (without Rif) and 4 h (with Rif). Rif-treated cells were washed three times with 500 µL complete MEM and incubated in the medium containing 50 µg/mL Gen at 37 °C for 2–3 h to allow for resuscitation of the intracellular *S. aureus* persisters. Infected cells were then treated with 2 µM aTc in the presence of 50 µg/mL Gen at 37 °C for 3 h to induce GFP and mKate expression. Macrophages harboring GFP-expressing *S. aureus* were imaged using a Leica SP8X Falcon (Leica Microsystems) with the following settings: (1) a tunable white light laser (WLL2) set to 484 nm was used for excitation with fluorescence detection at 495–570 nm and a detector gain of 100 V; (2) a 1024 X 1024 pixel scan at 16-bit with a scanning speed set to 600 was applied; (3) the pixel averaging was set to 2; (4) the pinhole size was set to 1 AU; (5) single images were obtained using a 63x oil-immersion objective lens (Plan-Apochromat, NA 1.4) with an 1 to 4-fold zoom. Macrophages infected with mKate-expressing *S. aureus* were stained with 100–300 nM LysoTracker Green DND-26 (ThermoFisher Scientific) at 37 °C for 30 min. The dye-loaded cells were washed once and kept in 200 µL complete MEM containing 1 µg/mL Hoechst 33342, 50 µg/mL Gen and 2 µM aTc. The z-stack acquisition and time-lapse live imaging were carried out using an Olympus FV3000RS confocal microscope (Olympus Life Science) equipped with a stage-top incubator at 37 °C, 5% CO_2_ and 100% humidity. The microscope settings were as follows: (1) 405-nm (Hoechst 33342), 488-nm (LysoTracker Green) and 594-nm (mKate) diode lasers were used for excitation with highly sensitive fluorescence spectral GaAsP detectors; (2) a 516 X 516 pixel scan was applied; (3) the pixel averaging was set to 10; (4) the pinhole size was set to 1 AU; (5) single images were obtained using a 60x oil-immersion objective lens (Super Correction, NA 1.4). All confocal images were exported as the default with no gamma correction by ImageJ/FIJI (62).

### Mouse infection and isolation of kidney cells

C57BL/6J mice (6- to 9-week-old) were systemically infected with the Tet-inducible GFP-expressing *S. aureus* strain HG003 (5×10^6^ CFU) via the tail vein intravenous route (15). At 1 day post-infection (dpi), mice were administered with 10 mg/kg Rif in 2.5% DMSO, 12.5% polyethylene glycol (PEG300) (Sigma) and 85% sterile H_2_O via intraperitoneal injection. Infected mice received the vehicle (2.5% DMSO/12.5% PEG300) were included as a control. At 2 dpi (24-h Rif treatment), dissected mouse kidneys were homogenized in 5 mL cold PBS containing deoxyribonuclease I (50 U/mL, bovine pancreas) (ThermoFisher Scientific) using a Stomacher 80 Biomaster (Seward, USA) at fast speed for 2 min twice (63). Tissue homogenates were filtered through 70-µm strainers (VWR International) and pelleted at 300 *g* for 5 min at 4 °C. After three washes in 4 mL cold PBS in 5-mL polypropylene round-bottom tubes (Corning) at 300 *g* for 5 min at 4 °C, cells were resuspended in cold PBS containing 1% fetal bovine serum and 50 µg/mL Gen in the presence or absence of 2 µM aTc for 3 h. The cell suspensions were probed with LIVE/DEAD fixable violet dead cell stain (1 µL/1 mL) (ThermoFisher Scientific) at 4 °C for 30 min before ImageStream analysis.

To evaluate the adjuvant activity of **KL1** in vivo, C57BL/6J mice were infected by intravenous injection with 2×10^7^ CFU of *S. aureus* strain HG003. At 6 hours post-infection (hpi), mice were administered with 10 mg/kg Rif, either alone or in combination with 100 mg/kg **KL1** in 7% DMSO, 40% PEG300 and 53% sterile H_2_O via intraperitoneal injection. Rif and **KL1** were administered once a day (q.d.) and every 12 h (b.i.d.), respectively, for 2 days. After 48 h treatment, dissected mouse organs were homogenized either by repetitively rolling a serological pipette over the samples in the sample bags (liver) or by bead beating using a Precellys 24 Touch (Bertin Technologies) at 5,000 rpm for 25 s twice with a 5 s interval (spleen). Tissue homogenates were 10-fold serially diluted in 1% NaCl and plated on TSA for the enumeration of survivor bacteria. The bacterial burden (CFU/g) was calculated by normalizing the number of CFU to the tissue weight.

### ImageStream analysis

Cell suspensions were loaded into an Amnis ImageStreamX Mark II (EMD Millipore) for single-cell imaging and fluorescence detection. For capturing live cells harboring viable GFP-expressing *S. aureus*, 405-nm and 488-nm colinear lasers were set to 40 mW and 70–90 mW for the live/dead cell discrimination and GFP expression, respectively. In-focus single-cell images were acquired with a 60X magnification and a low-speed flow rate using the INSPIRE software (EMD Millipore). The exported data were analyzed using IDEAS 6.2 software (EMD Millipore).

### High-throughput screen for intracellular *S. aureus* energy modulators

RAW 264.7 macrophages were seeded in 4 mL complete MEM in Costar 6-well tissue culture-treated plates (Corning) (8×10^5^ cells/well) at 37 °C and 5% CO_2_ for 16 h. Cells were infected with *S. aureus* strain JE2–Lux at an MOI of 100 at 37 °C for 25 min. The infected macrophages were washed once with 2 mL PBS and incubated with 10 mM EDTA in PBS (2 mL/well) at 37 °C for 5–10 min. After dissociation, cells were pelleted at 300 *g* for 6 min, resuspended in complete MEM containing 50 µg/mL Gen at a density of 2.5×10^5^ cells/mL and dispensed 80 µL of the cell suspension into each well of 384-well tissue culture-treated white plates and optically clear polymer bottom black plates (ThermoFisher Scientific) for luminescence and fluorescence detections, respectively. Uninfected macrophages were also added to column 1 and 24 of each plates as controls for the luminescence background and the baseline of host cell viability. The plates were preloaded with 80 nL of 10 mM compounds using a Mosquito LV multichannel pipetting system (SPT Labtech, Covina). The vehicle control (0.1% DMSO) and 10 µg/mL Rif were included in each experiment for comparison. After cell dispensing, the plates were centrifuged at 500 *g* for 2 min and incubated in a humidified chamber at 37 °C and 5% CO_2_ for 4 h. The luminescence signals were measured at an 1.25-mm depth with a detector gain of 250 at 37 °C using a Synergy H1 microplate reader. To monitor the cell viabilities after compound treatment, 10 µL CellTiter-Fluor reagent (Promega) were added to the black plates and incubated in the dark at 37 °C and 5% CO_2_ for 30 min. The fluorescence intensities were measured at a 7-mm depth with a detector gain of 100 at 37 °C. The kinase-targeted compound library containing >4,700 rule of five-compliant compounds were part of the chemical collection at the Center for Integrative Chemical Biology and Drug Discovery (CICBDD) at the University of North Carolina at Chapel Hill. The luminescence and fluorescence signals were normalized to the corresponding vehicle controls (0.1% DMSO). EC_50_ values of Van and Rif for the intracellular MRSA strain JE2–Lux were determined by fitting the luminescence data to a standard dose response equation (GraphPad Prism).

### Reactive species assays

The reactive oxygen and nitrogen species were measured using a luminescent probe L-012 (Wako Chemical Corporation) and a fluorescent dye fluorescein-boronate (Fl-B) (56). For L-012 probing, RAW 264.7 macrophages were seeded in 100 µL complete MEM in Falcon 96-well tissue culture-treated white plates (Corning) (3.84×10^4^ cells/well) at 37 °C and 5% CO_2_ for 16 h. Cells were infected with *S. aureus* strain JE2 at an MOI of 10 by centrifugation at 1,000 *g* for 2 min, followed by incubation at 37 °C for 40 min. Cells were treated with 40–100 µM **KL1** or 1–10 µM BIX-01294 in the presence of 50 µg/mL Gen in five replicates at 37 °C and 5% CO_2_ for 4–8 h. After three washes with 200 µL pre-warmed PBS, 100 µL of 300 µM pre-warmed Hanks’ balanced salt solution (Gibco) was added to each well. The luminescence signals were immediately measured with auto-gain at 37 °C using a Synergy H1 microplate reader. The signals of two consecutive reads were averaged to minimize the technical variability between reads. For Fl-B staining, macrophages were seeded in 96-well tissue culture-treated black clear-bottom plates (Corning) with the same culture condition. After treatment with 40–100 µM **KL1**, 100 µM KL2, 40–100 µM the inactive analog KL7, 1–10 µM BIX-01294 or 20 µM butylated hydroxyanisole (Sigma), infected cells were washed twice with 200 µL pre-warmed PBS and incubated with 100 µL of 25 µM Fl-B in PBS at 37 °C for 30 min. Cells were washed three times with 200 µL PBS to remove unbound dyes and maintained in 100 µL PBS. The fluorescence signals were measured at 535 nm with excitation at 485 nm with a detector gain of 100 and an area scan mode using a Synergy H1 microplate reader.

### ATP measurement

*S. aureus* strains LAC, HG003 and JE2 were grown in TSB for 16–18 h and subcultured at 1:1,000 to 1:40,000 dilution in fresh TSB containing 40 µM **KL1** or 0.1–0.5% DMSO in six replicates at 37 °C for 4 h. The bacterial cultures were then aliquoted into Falcon 96-well white plates (100 µL/well) and mixed with one volume of BacTiter-Glo reagent (Promega), followed by incubation in the dark on a rocker with gentle shaking at room temperature for 20 min. The luminescence signals were detected at a 1.25-mm depth with a detector gain of 250 using a Synergy H1 microplate reader and normalized to the corresponding CFUs to determine the relative ATP levels.

### Seahorse analysis

The stationary-phase bacterial cultures were diluted at 1:100 in fresh TSB in triplicate and incubated for 2–3 h until the OD_600_ values reached 0.3. Each culture was then diluted at 1:500 to 1:1,000 in TSB or Seahorse XF DMEM (Agilent Technologies) supplemented with 100 mM glucose, 2 mM L-glutamine and 1 mM sodium pyruvate and dispensed into a poly-D-lysine (PDL)-coated Seahorse XF HS miniplate (100 µL/well). PDL-coating was performed by adding 100 µg/mL PDL in sterile H_2_O to each well of the miniplate for 30 min. Excessive PDL was removed by two washes with H_2_O, and the miniplate was air-dried prior to the assays. Bacteria adherence was achieved by centrifugation at 1,400 *g* for 10 min. An additional 80 µL media was carefully added to each well (180 µL final volume), and the miniplate was loaded into a pre-calibrated Seahorse XF HS Mini Analyzer (Agilent Technologies). A final concentration of 40 µM of **KL1** or 0.1–0.5% DMSO was injected into each well, and the oxygen consumption rates (OCRs) and extracellular acidification rates (ECARs) of the bacteria were monitored at 37 °C for 4–6 h. The data was analyzed using Seahorse Analytics (Agilent Technologies) and plotted using GraphPad Prism.

### Transcriptomic analysis

RAW 264.7 macrophages were seeded in 4 mL complete MEM in 6-well tissue culture-treated plates (8×10^5^ cells/well) at 37 °C and 5% CO_2_ for 16 h. Cells were infected with

*S. aureus* strain JE2–Lux at an MOI of 20 at 37 °C for 35 min and incubated in fresh media containing 100 µg/mL Gen and 40 µM **KL1** or 0.1% DMSO at 37 °C for 24 h. After a PBS wash, cells were dissociated with 10 mM EDTA in PBS (2 mL/well) at 37 °C for 5 min, followed by three washes in 10 mL PBS at 4 °C to remove EDTA. The number of viable host cells was enumerated to ensure consistent sample input (2.4–4.2×10^6^ total viable cells for both **KL1**- and DMSO-treated groups). The frozen cell pellets were delivered to Azenta Life Sciences for the standard RNA-sequencing (paired-end, 30 million reads per sample) and data analysis. The aligned sequencing data were mapped to mouse GRCm38 reference genome available on ENSEMBL using the STAR aligner v.2.5.2b.

### Databases

Biological test results of the identified compound **KL1** (PubChem CID: 2881454) were retrieved from PubChem (https://pubchem.ncbi.nlm.nih.gov/) (64). Gene expression profiles were examined using Expression Atlas (https://www.ebi.ac.uk/gxa/home) (65). Biological functions and subcellular localization data were sourced from the UniProt Knowledgebase (https://www.uniprot.org/) (37). Protein association networks among the differentially expressed genes were analyzed using STRING (https://string-db.org/) (66). Chemical structures were gene rated using ChemDraw software 21.0.0 (PerkinElmer).

## Acknowledgements

This work was supported by the Burroughs Wellcome Fund (BWF) in Pathogenesis of Infectious Disease (PATH award to B.P.C), the National Institutes of Health (NIH) (R01AI173004 and R01AI179695 to B.P.C and R01AI167978 to S.E.R) and UNC Lineberger Comprehensive Cancer Center. The content of this study is solely the responsibility of the authors and does not necessarily represent the official views of the funding agents. We are grateful to the UNC Neuroscience Microscopy Core and the UNC Flow Cytometry Core Facility (supported in part by P30 CA016086). Part of the figures was created with BioRender.com.

## Contributions

K.L. and B.P.C. conceptualized the project and wrote the manuscript. K.L. performed the genetic engineering, in vitro and animal experiments. K.L. and N.J.W. performed the Seahorse analyses. K.L., B.H. and M.A. performed the high-throughput screen. X.Y. synthesized the compounds. M.J.G.E. performed the *Salmonella* infection assays. V.G.F. provided *S. aureus* clinical isolates. B.P.C. and S.E.R. provided funding for the project. B.P.C., S.E.R., X.W., S.H. and K.H.P. supervised the study. All the authors reviewed and edited the manuscript.

## Footnotes

To whom correspondence should be addressed. The authors have no competing interests.

---

**Supplementary table 1.**
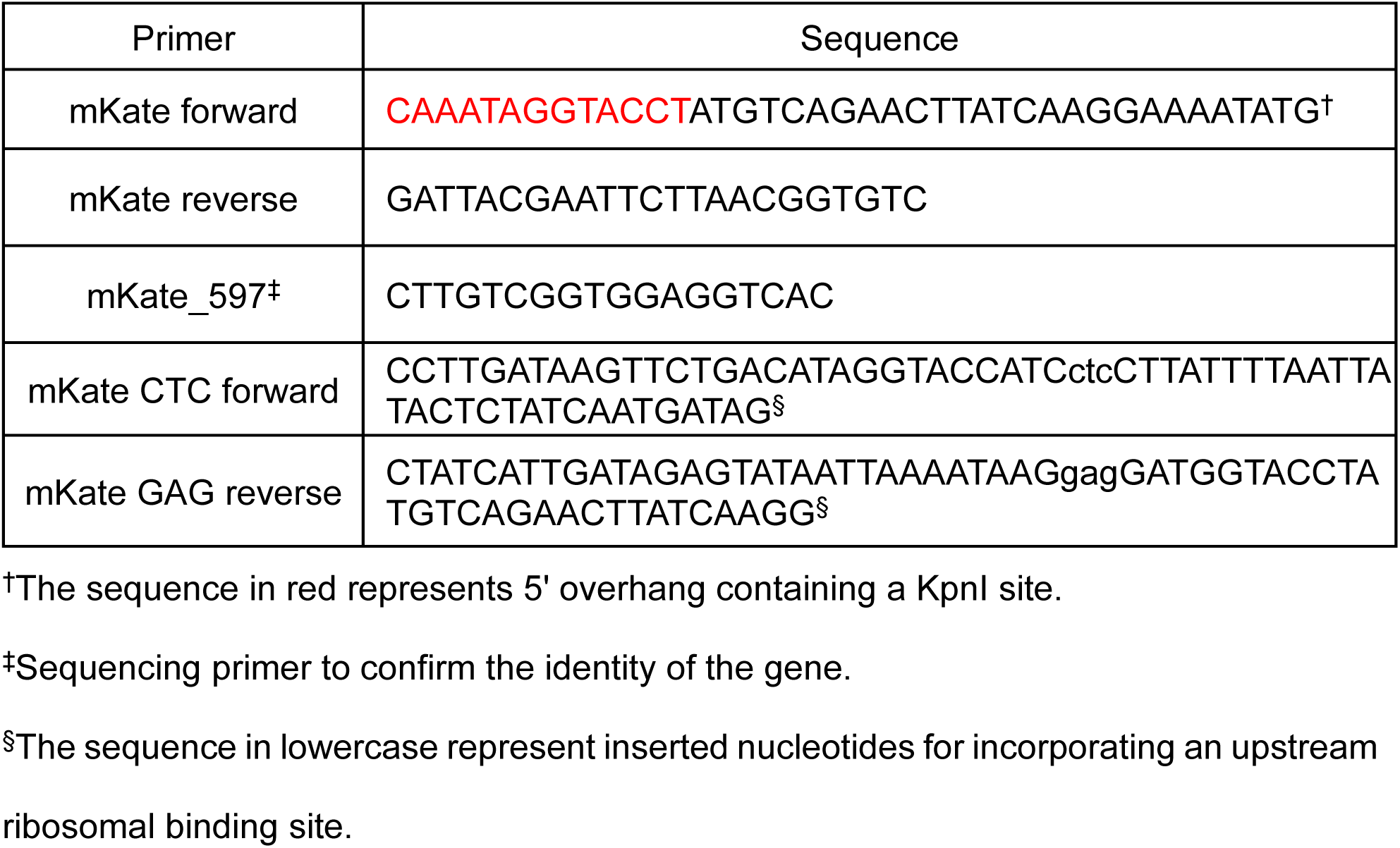
Primers for molecular cloning.

**Supplementary table 2.**
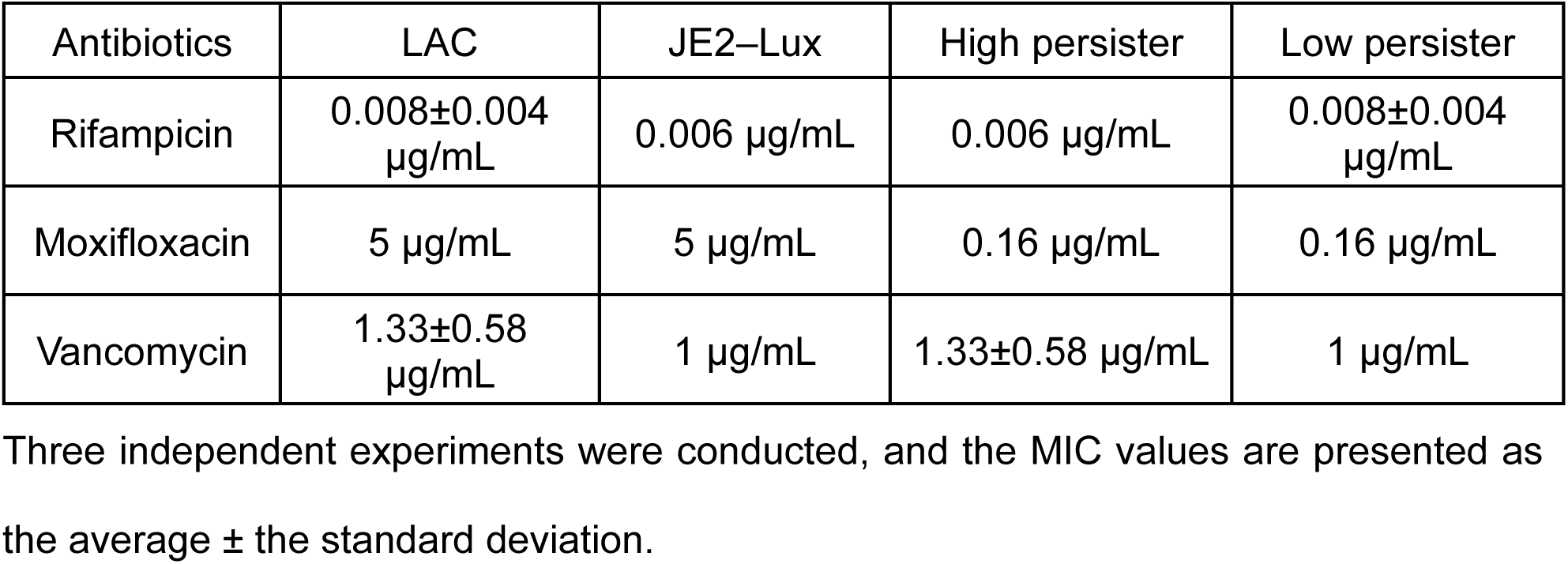
Minimum inhibitory concentrations (MICs) of antibiotics in the *S. aureus* strains used in this study.

**Supplementary table 3.**
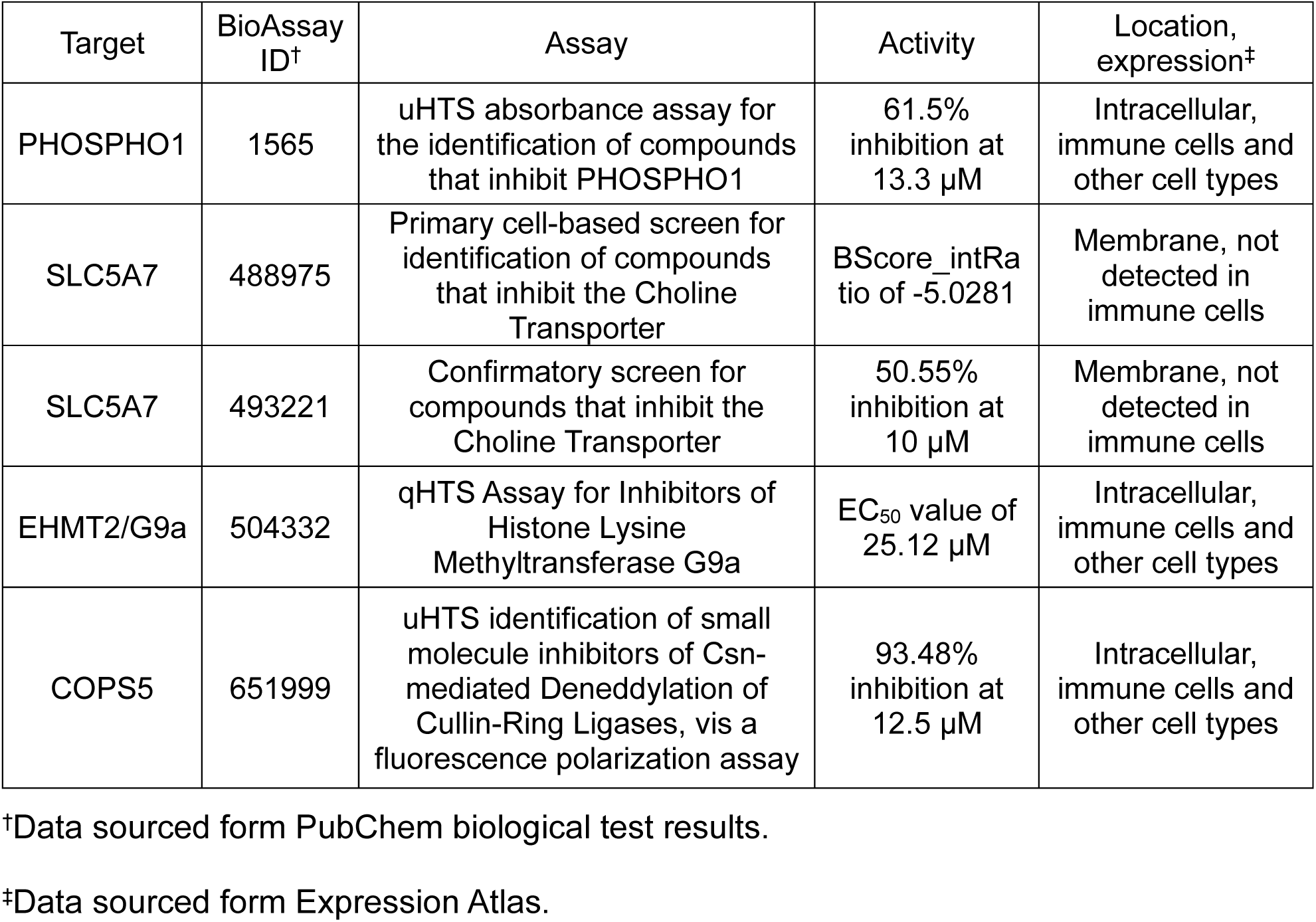
Potential targets of KL1 based on published functional screens.

| Target | BioAssay ID<sup>†</sup> | Assay | Activity | Location, expression<sup>‡</sup> |
| --- | --- | --- | --- | --- |
| PHOSPHO1 | 1565 | uHTS absorbance assay for the identification of compounds that inhibit PHOSPHO1 | 61.5% inhibition at 13.3 μM | Intracellular, immune cells and other cell types |
| SLC5A7 | 488975 | Primary cell-based screen for identification of compounds that inhibit the Choline Transporter | BScore_intRatio of -5.0281 | Membrane, not detected in immune cells |
| SLC5A7 | 493221 | Confirmatory screen for compounds that inhibit the Choline Transporter | 50.55% inhibition at 10 μM | Membrane, not detected in immune cells |
| EHMT2/G9a | 504332 | qHTS Assay for Inhibitors of Histone Lysine Methyltransferase G9a | EC<sub>50</sub> value of 25.12 μM | Intracellular, immune cells and other cell types |
| COPS5 | 651999 | uHTS identification of small molecule inhibitors of Csn-mediated Deneddylation of Cullin-Ring Ligases, vis a fluorescence polarization assay | 93.48% inhibition at 12.5 μM | Intracellular, immune cells and other cell types |

<sup>†</sup>Data sourced form PubChem biological test results.
<sup>‡</sup>Data sourced form Expression Atlas.

**Supplementary figure 1.**
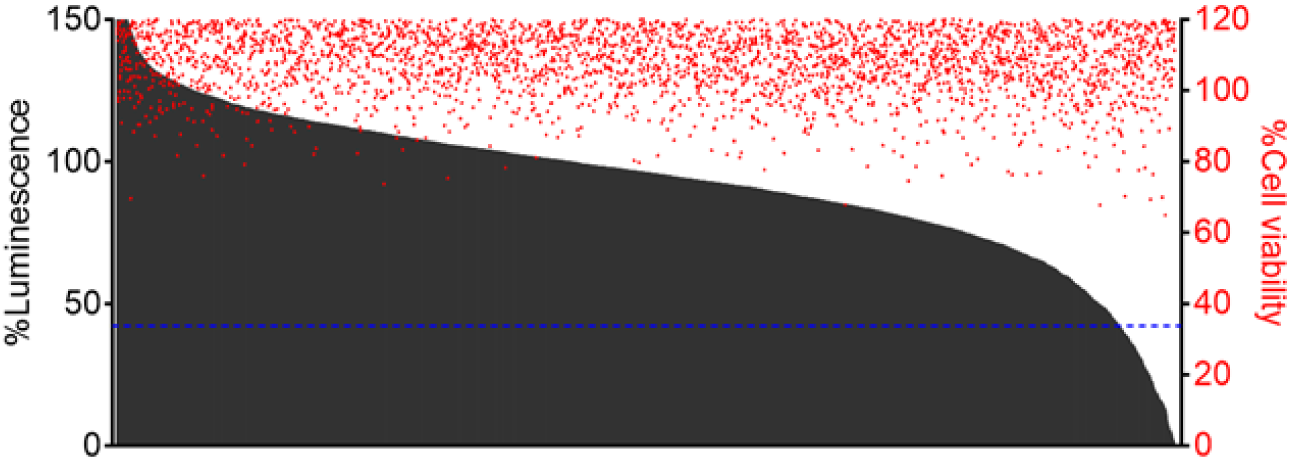
High-throughput screening of a kinase-targeted library identifies compounds with differential capacities in modulating the energy state of intracellular *S. aureus*. A total of >4,700 compounds (10 µM) from the UNC CICBDD were screened for targeting the intracellular MRSA. All of these compounds shared structural similarity to kinase inhibitors and complied with the Lipinski’s rule of five. The luminescence signals were normalized to the vehicle controls (0.1% DMSO) (black bars). The normalized host cell viability was measured by the CellTiter-Fluor assay (red circles). The %luminescence of 10 µg/mL rifampicin-treated cells is indicated as a reference (blue dashed line).

**Supplementary figure 2.**
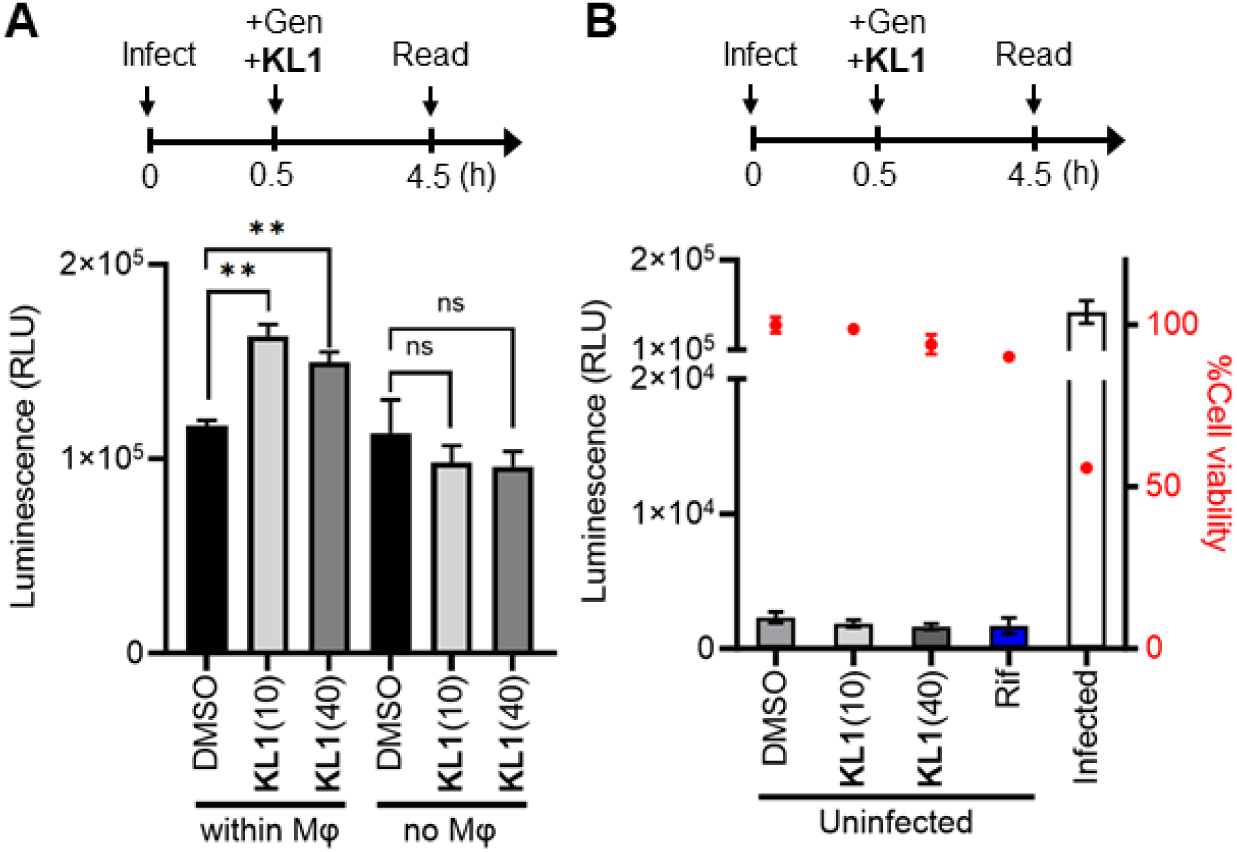
The energy state up-regulation by KL1 is not caused by background noise from the compound, the host cells or dead extracellular bacteria. (**A**) Macrophages infected with the bioluminescent MRSA strain JE2–lux (within Mφ) were treated with 10 µM or 40 µM **KL1**. Gentamicin (Gen; 50 µg/mL) was added to eliminate the extracellular bacteria. The bacterial cultures without macrophages (no Mφ) in the presence of Gen were examined with the same input CFU and culture conditions (n = 3). (**B**) Uninfected macrophages were treated with 10 µM or 40 µM **KL1**. The normalized host cell viability was measured by the CellTiter-Fluor assay (red circles). Rifampicin (Rif)-treated cells (blue bar) and infected macrophages (white bar) were included as controls (n = 3). Assay schematics are shown above plots. **p<0.01; ns, not significant (unpaired t-test). The bars represent mean ± SEM.

**Supplementary figure 3.**
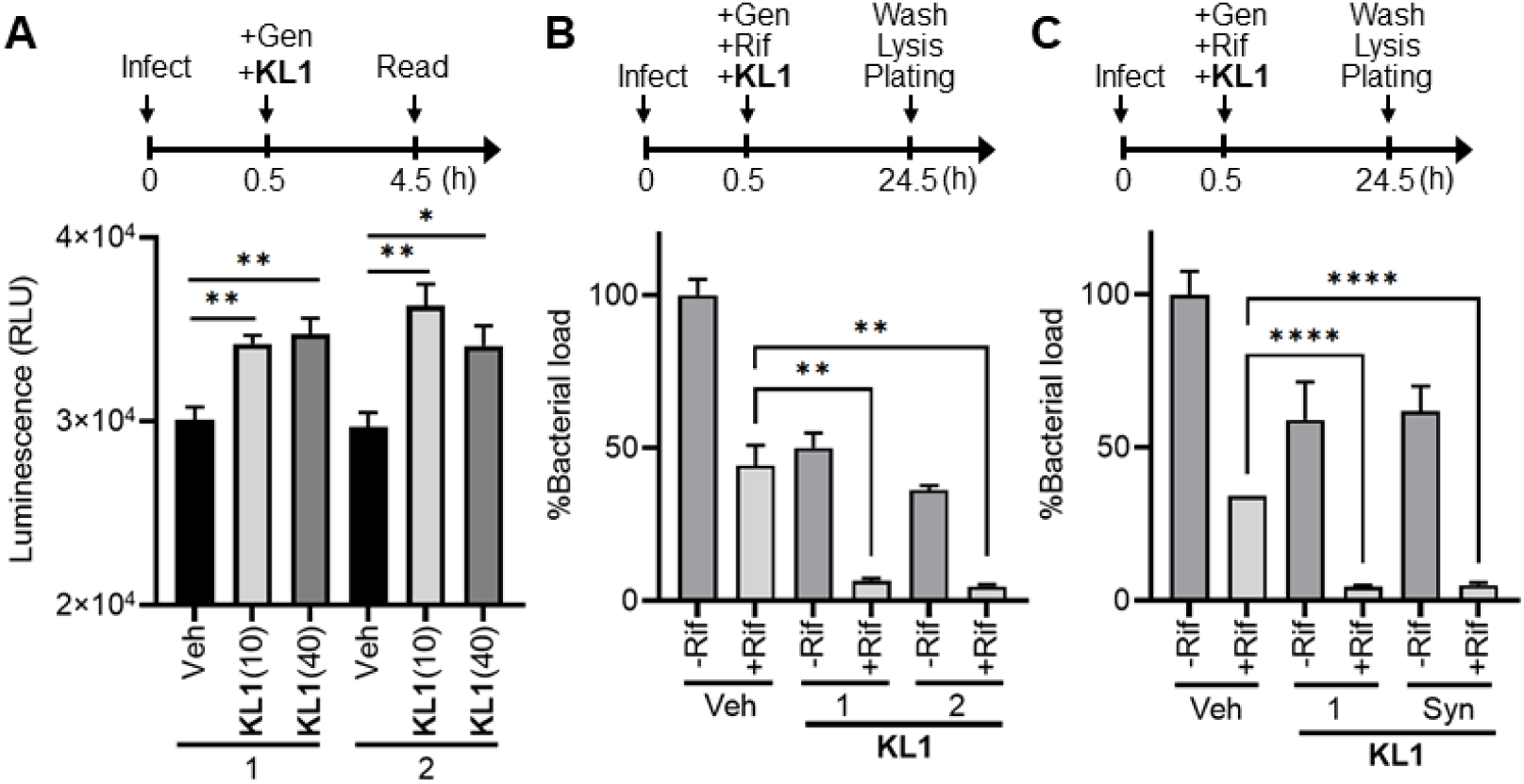
The adjuvant activity of KL1 was consistently observed with the compound across different sources. (**A** and **B**) **KL1** (10 and 40 µM) purchased from two independent suppliers, ChemBridge (1) and Enamine (2), consistently elevated the energy state of intracellular *S. aureus* (n = 4) and sensitized the bacteria to killing by rifampicin (Rif, 10 µg/mL) (n = 3). (**C**) **KL1** synthesized in-house (Syn) displayed adjuvant activity comparable to that of commercially available **KL1** (n = 3). Gentamicin (Gen; 50 µg/mL) was added to eradicate the extracellular bacteria. The numbers of tolerant bacteria were normalized to the untreated control (no Rif, no **KL1**, Gen-only). Assay schematics are shown above plots. *p<0.05; **p<0.01; ****p<0.0001 (unpaired t-test). The bars represent mean ± SEM.

**Supplementary figure 4.**
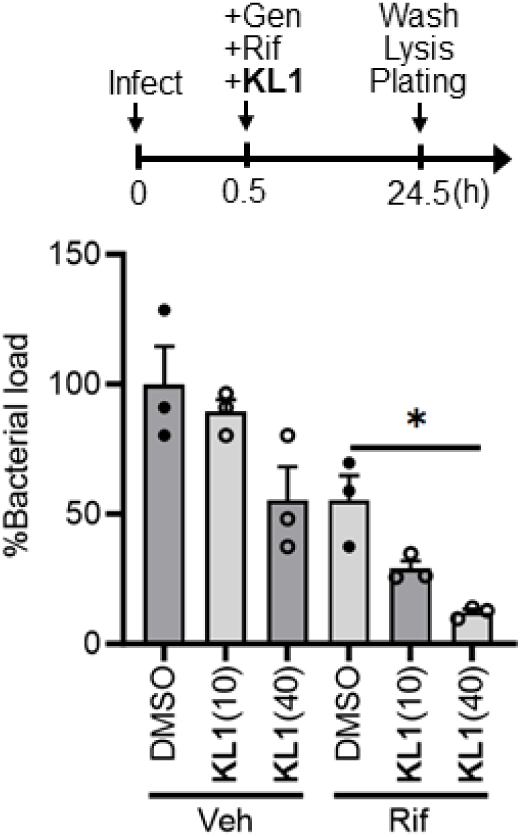
Compound KL1 sensitizes the killing of intracellular MRSA by antibiotics in a dose-dependent manner. RAW 264.7 cells were infected with the MRSA strain JE2–lux and treated with 10 µM or 40 µM **KL1** in the presence or absence of 10 µg/mL rifampicin (Rif). Gentamicin (Gen; 50 µg/mL) was added to eliminate the extracellular bacteria. The numbers of survivor bacteria were normalized to the untreated control (no Rif, no **KL1**, Gen-only) (n = 3). *p<0.05 (unpaired t-test). The bars represent mean ± SEM.

**Supplementary figure 5.**
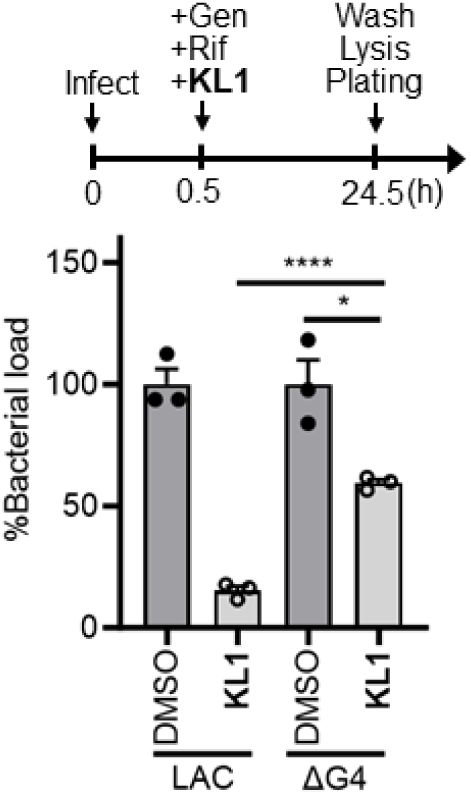
Glucose uptake is important for the synergism between KL1 and antibiotics. RAW 264.7 cells were infected with the wild-type strain LAC or the mutant lacking all four glucose transporters (ΔG4). The infected cells were treated with 40 µM **KL1** and 10 µg/mL rifampicin (Rif). Gentamicin (Gen) was included to eliminate the extracellular bacteria. The numbers of survivor bacteria were normalized to the corresponding controls (no **KL1**, Gen- and Rif-only) (black circles) (n = 3). *p<0.05; ****p<0.0001 (unpaired t-test). The bars represent mean ± SEM.

**Supplementary figure 6.**
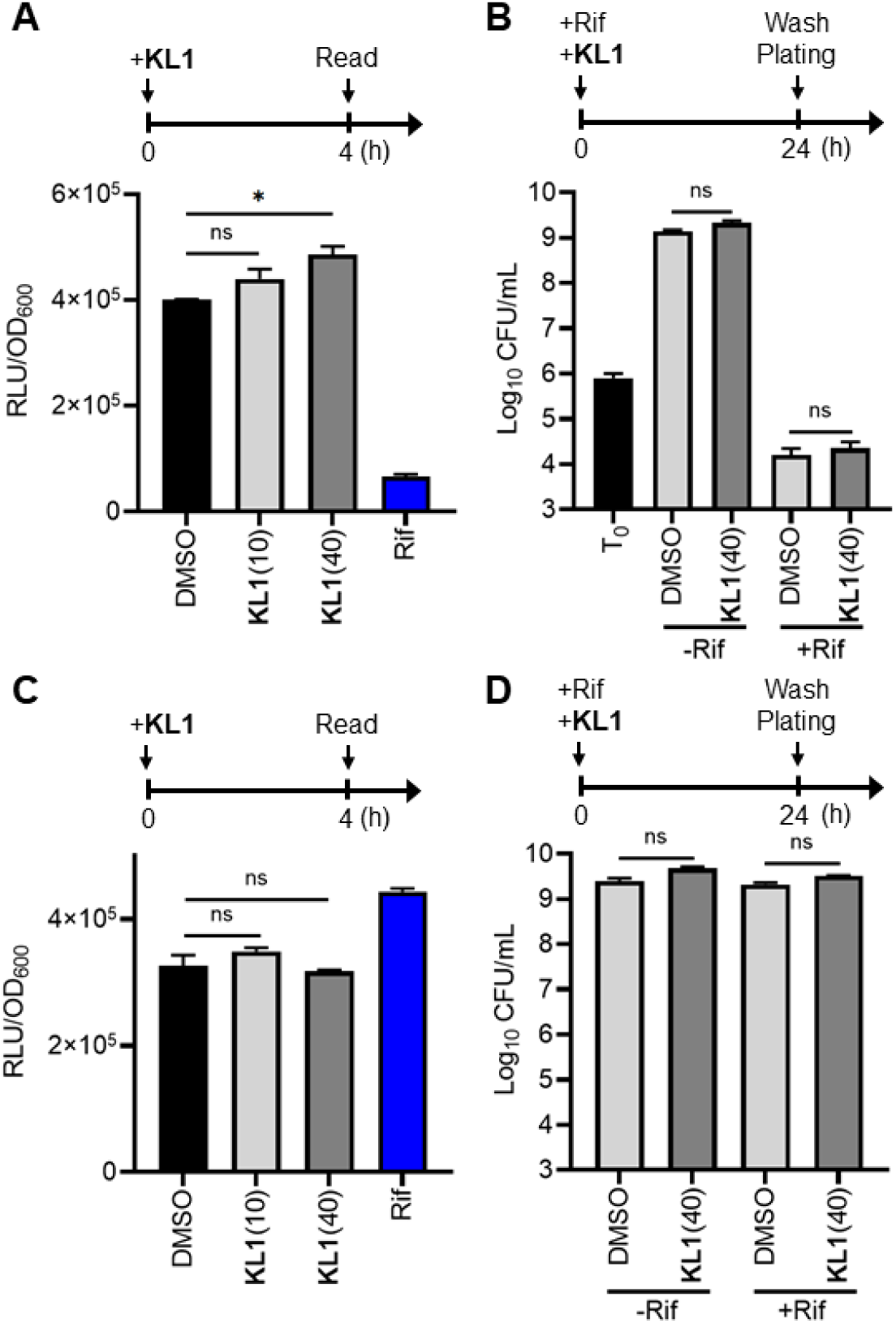
Compound KL1 exhibits a minute effect on the extracellular *S. aureus* activity and does not synergize with antibiotics in killing the extracellular bacteria. (**A** and **C**) The exponential-phase (**A**) and stationary-phase (**C**) *S. aureus* cultures (JE2–lux) were incubated with 10 µM or 40 µM **KL1** for 4 h, followed by luminescence detection. The signal intensities (RLU) were normalized to the OD_600_ values. The vehicle control (0.1% DMSO, black bar) and rifampicin (Rif; 2 µg/mL)-treated group (blue bar) were included as references. (**B** and **D**) The exponential-phase (**B**) and stationary-phase (**D**) cultures were treated with and without 40 µM **KL1** and 2 µg/mL Rif for 24 h, followed by a wash step and plating to enumerate the number of tolerant bacteria. Assay schematics are shown above plots. *p<0.05; ns, not significant (unpaired t-test). The bars represent mean ± SEM.

**Supplementary figure 7.**
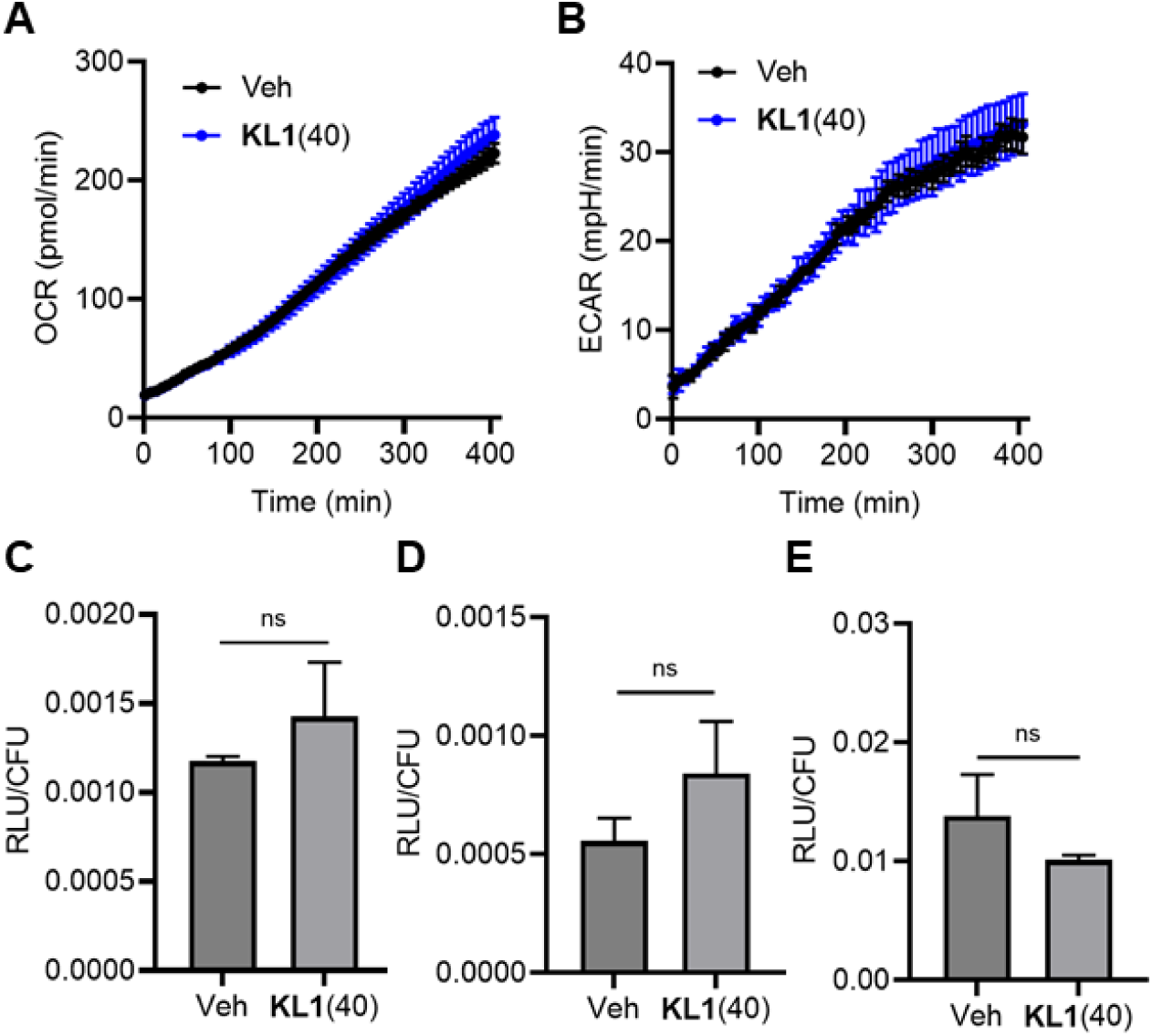
Compound KL1 does not impact the metabolic activity and ATP levels in extracellular bacteria. (**A** and **B**) Metabolic analysis indicates that **KL1** (40 µM, blue) does not alter the oxygen consumption rate (OCR) (**A**) and extracellular acidification rate (ECAR) (**B**) in extracellular *S. aureus* comparing to the vehicle control (Veh, 0.1–0.5% DMSO, black). Representative data of six independent experiments (n = 3) is shown. The bars represent mean ± SD. (**C**–**E**) Exponential-phase cultures were treated with 40 µM **KL1** or 0.1% DMSO for 4 h, followed by measuring the relative ATP levels using a BacTiter-Glo assay. The luminescent signals were normalized to the numbers of bacteria. *S. aureus* strains HG003 (**C**), JE2 (**D**) and LAC (**E**) were examined (n = 3–6). ns, not significant (unpaired t-test).

**Supplementary figure 8.**
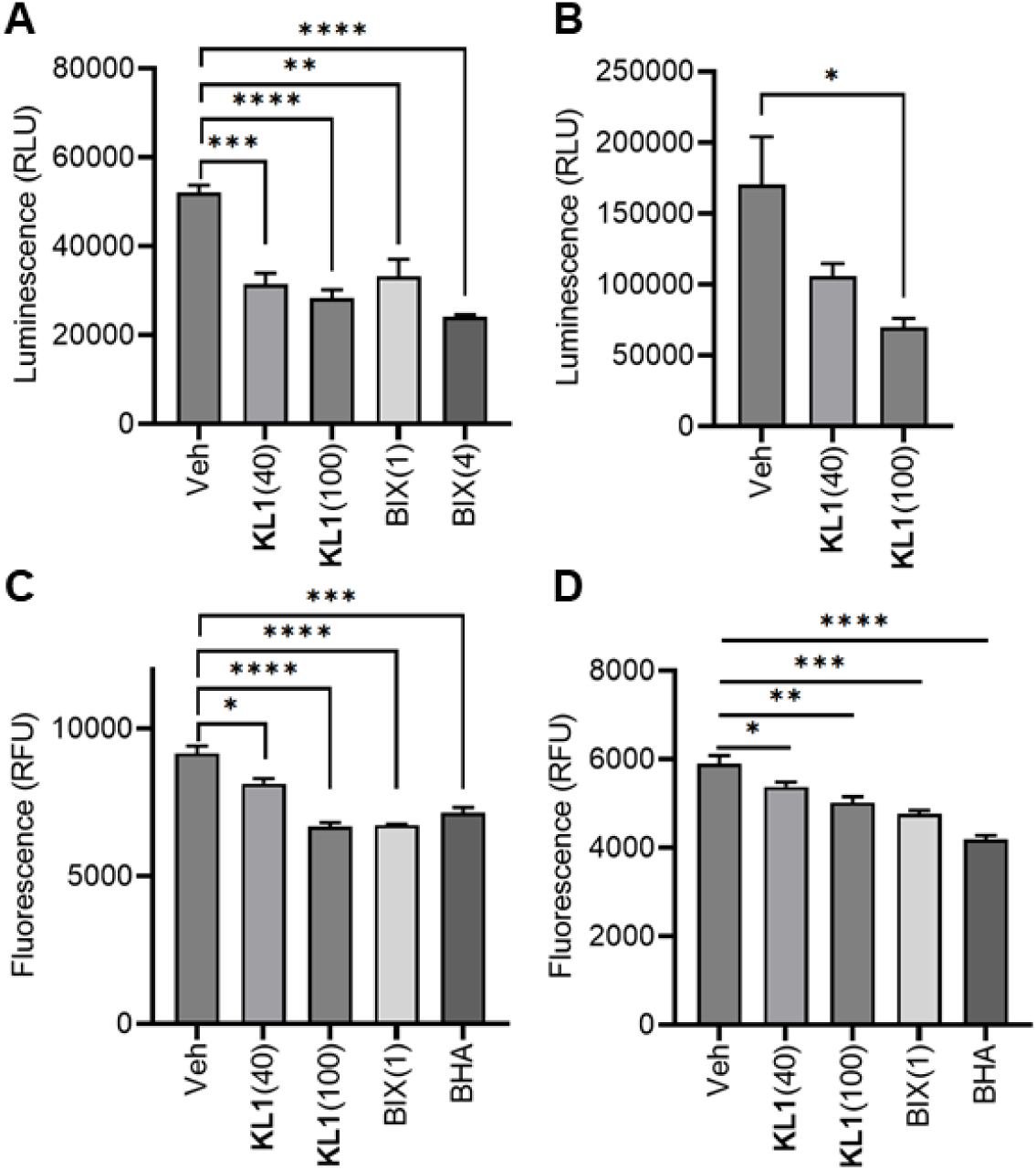
KL1 reduces the level of reactive species in both infected and uninfected macrophages. Compound **KL1** (40 and 100 µM) and the EHMT2/G9a inhibitor, BIX-01294 (BIX; 1 and 4 µM) decreased the ROS/RNS level in *S. aureus*-infected macrophages at 8 hours post-infection (hpi) (**A** and **C**) and in uninfected macrophages (**B** and **D**). L-012 (**A** and **B**) and fluorescein-boronate (Fl-B) (**C** and **D**) were used to quantify the ROS/RNS level. DMSO (Veh; 0.25%) and butylated hydroxyanisole (BHA; 20 µM) were included as controls. Representative data of three independent experiments are shown (n = 5). *p<0.05; **p<0.01; ***p<0.001; ****p<0.0001 (unpaired t-test). The bars represent mean ± SEM.

**Supplementary figure 9.**
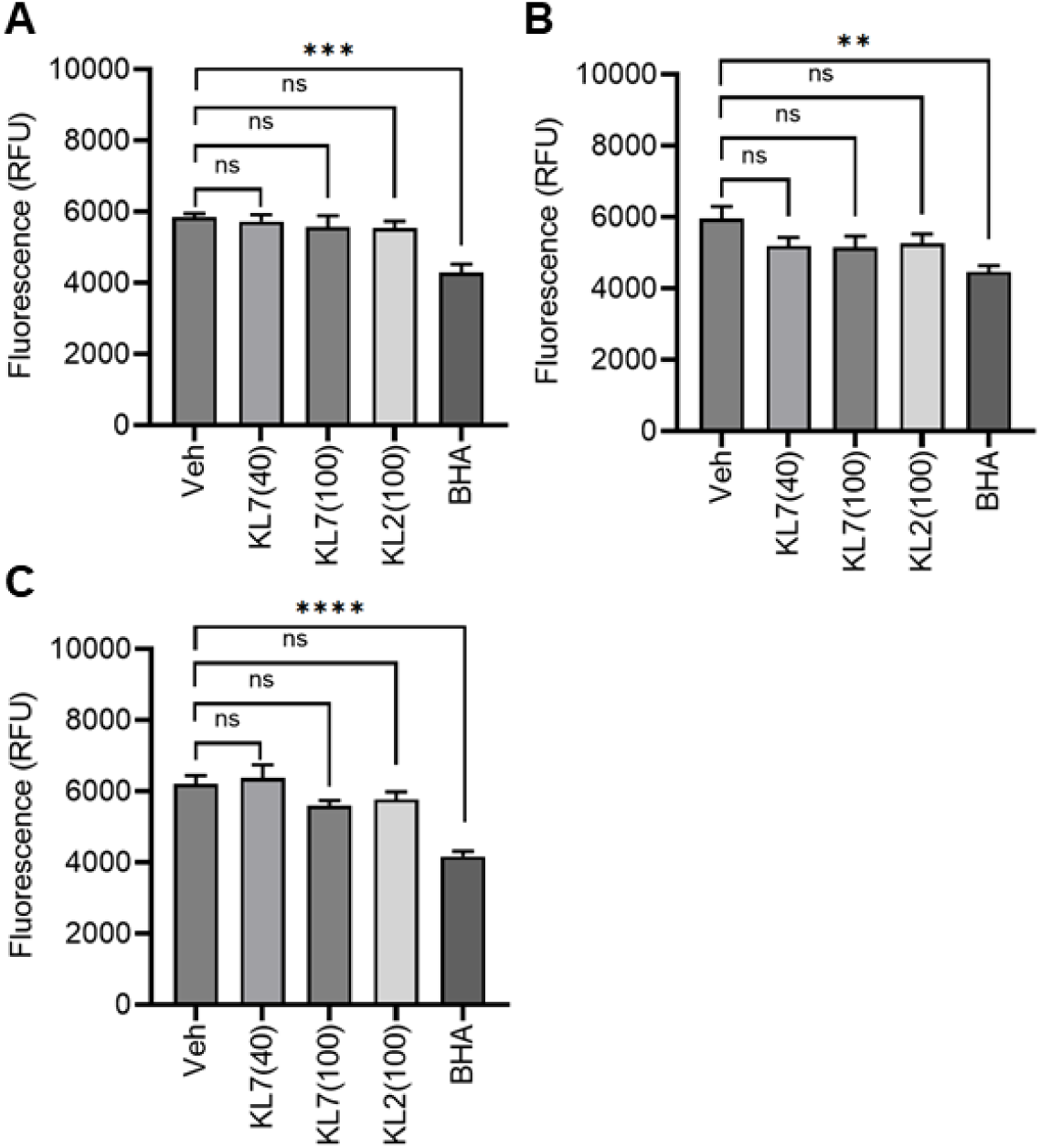
The inactive analog of KL1 losses the ability to reduce the level of reactive species in both infected and uninfected macrophages. The inactive analog KL7 (40 and 100 µM) and analog KL2 (100 µM) with lower adjuvant activity did not affect ROS/RNS production in uninfected (**A** and **B**) and *S. aureus*-infected macrophages (**C**) after 4 h (**A**) and 8 h (**B** and **C**) treatment. Fluorescein-boronate (Fl-B) was used to quantify the level of reactive species. DMSO (Veh; 0.25%) and butylated hydroxyanisole (BHA; 20 µM) were included as controls. Representative data of three independent experiments are shown (n = 5). **p<0.01; ***p<0.001; ****p<0.0001; ns, not significant (unpaired t-test). The bars represent mean ± SEM.

