## Supplementary Information for "A host-directed adjuvant resuscitates and sensitizes intracellular bacterial persisters to antibiotics"

**This PDF file includes:**

- Supplementary text
- Figures S1 to S9
- Table S1 to S3
- Legends for Movies S1 to S3

**Other supplementary materials for this manuscript include the following:**

- Movies S1 to S3

### Supplementary Information Text

### Chemical synthesis of KL1 and KL7

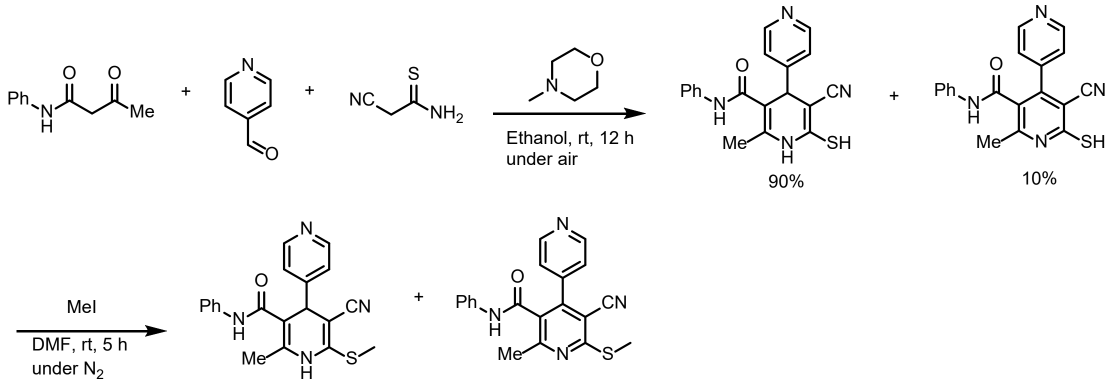


**KL-1** **KL-7**

4-Methylmorpholine (3.29 mL, 30 mmol) was added dropwise to a solution of 3-oxo-N-phenylbutanamide (3.54 g, 20 mmol), isonicotinaldehyde (2.14 g, 20 mmol) and 2-cyanoethanethioamide (2.00 g, 20 mmol) in 150 mL ethanol. The reaction mixture was stirred at room temperature for 12 h under air atmosphere. Afterwards, volatiles were removed, yielding a viscous oil, which was redissolved in 20 mL dichloromethane (DCM). Then, 150 mL of hexane was added to the solution. The mixture was stirred for an additional 10 min at room temperature, resulting in a slurry. The slurry was filtered, and the solid residue was washed with a mixture of DCM/hexane (1:15, v/v). The product was dried under high vacuum to give 5-cyano-6-mercapto-2-methyl-N-phenyl-1,4-dihydro-[4,4'-bipyridine]-3-carboxamide with >90% purity (4.02 g, 10.40 mmol).

**MS (ESI)**: m/z calculated for $C_{19}H_{17}N_4OS$: 349.11 $[M + H]^+$; found 349.10.

100 mg of the crude product was further purified by normal-phase ISCO chromatography. The isolated impurity was isolated and characterized as the oxidized analog 5-cyano-6-mercapto-2-methyl-N-phenyl-[4,4'-bipyridine]-3-carboxamide, as the major

byproduct.

**MS (ESI)**: m/z calculated for $C_{19}H_{15}N_4OS$: 347.10 $[M + H]^+$; found 347.10.

**$^1$H NMR** (400 MHz, DMSO-*$d_6$*) δ 14.52 (s, 1H), 10.33 (s, 1H), 8.71–8.65 (m, 2H), 7.48–7.42 (m, 2H), 7.34–7.20 (m, 4H), 7.10–7.01 (m, 1H), 2.49 (s, 3H).

For methylation, 538 μL methyl iodide (MeI, 8.61 mmol) dissolved in 3 mL dimethylformamide (DMF) was added dropwise to a solution of the crude 5-cyano-6-mercapto-2-methyl-N-phenyl-1,4-dihydro-[4,4'-bipyridine]-3-carboxamide (3.33 g, 8.61 mmol) in 50 mL DMF. The mixture was stirred at room temperature for 5 h under nitrogen atmosphere. Water was then added, and the mixture was extracted three times with ethyl acetate (EA). The organic phases were combined, washed with brine, dried over sodium sulfate and filtered. Volatiles were removed, and the residue was purified by normal-phase ISCO chromatography. The crude product was further slurried with a mixture of DCM/hexane (1:15, v/v), affording 5-cyano-2-methyl-6-(methylthio)-N-phenyl-1,4-dihydro-[4,4'-bipyridine]-3-carboxamide (1.98 g, 5.46 mmol) (**KL1**) with >99% purity.

**MS (ESI)**: m/z calculated for $C_{20}H_{19}N_4OS$: 363.13 $[M + H]^+$; found 363.20.

**$^1$H NMR** (400 MHz, DMSO-*$d_6$*) δ 9.73 (s, 1H), 9.18 (s, 1H), 8.56–8.50 (m, 2H), 7.56–7.48 (m, 2H), 7.29–7.22 (m, 2H), 7.22–7.18 (m, 2H), 7.01 (tt, *J* = 7.2, 1.2 Hz, 1H), 4.75 (s, 1H), 2.52 (s, 3H), 2.13–2.08 (m, 3H).

**$^{13}$C NMR** (100 MHz, DMSO-*$d_6$*) δ 165.95, 152.22, 150.05, 147.04, 138.94, 137.19, 128.56, 123.33, 122.19, 119.64, 119.35, 106.12, 83.12, 42.30, 17.07, 15.63.

Fractions containing the oxidized impurity were also collected (KL7). Volatiles were removed to yield 5-cyano-2-methyl-6-(methylthio)-N-phenyl-[4,4'-bipyridine]-3-carboxamide as a white solid in 98% purity.

**MS (ESI)**: m/z calculated for $C_{20}H_{17}N_4OS$: 361.11 $[M + H]^+$; found 361.10.

**$^1$H NMR** (400 MHz, DMSO-$d_6$) δ 10.45 (s, 1H), 8.74–8.65 (m, 2H), 7.51–7.45 (m, 2H), 7.40–7.32 (m, 2H), 7.32–7.22 (m, 2H), 7.12–7.03 (m, 1H), 2.70 (s, 3H), 2.64 (s, 3H).

**$^{13}$C NMR** (100 MHz, DMSO-$d_6$) δ 163.26, 162.54, 158.59, 149.81, 148.48, 141.85, 137.87, 128.84, 128.03, 124.35, 122.98, 119.62, 114.48, 103.13, 23.04, 13.00.

### $^1$H NMR Spectrum (KL1)

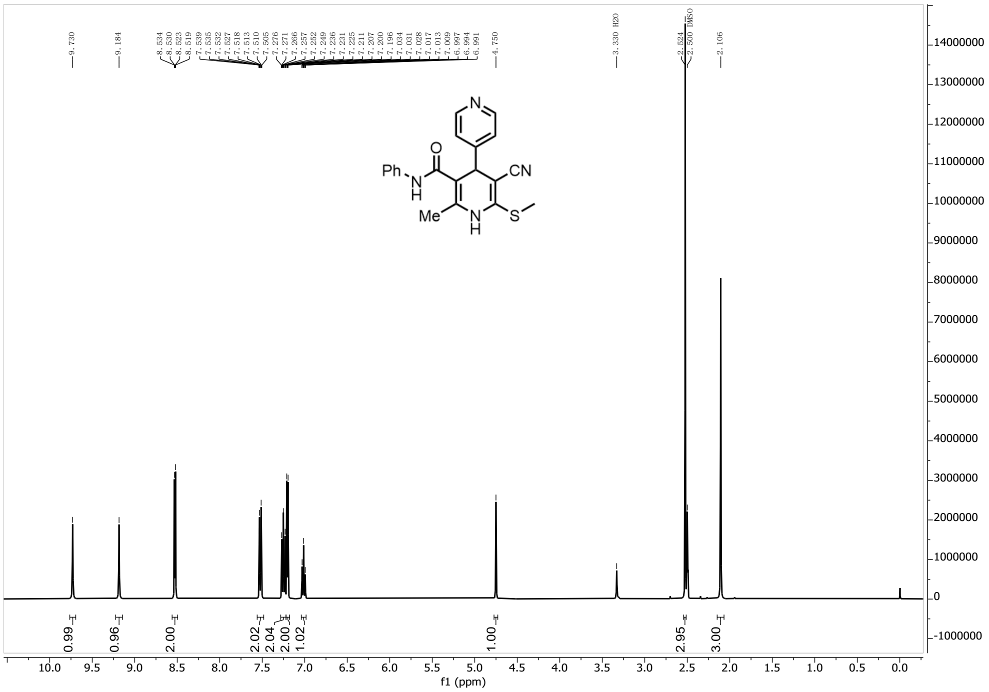


### $^1$H NMR Spectrum (KL7)

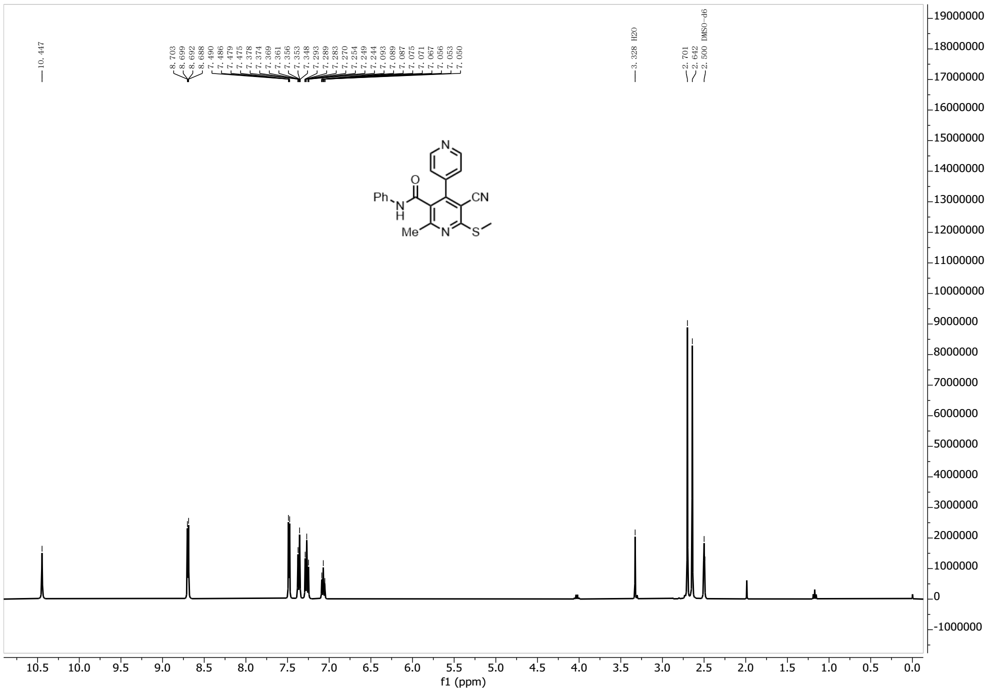


### $^{13}C$ NMR Spectrum (KL1)

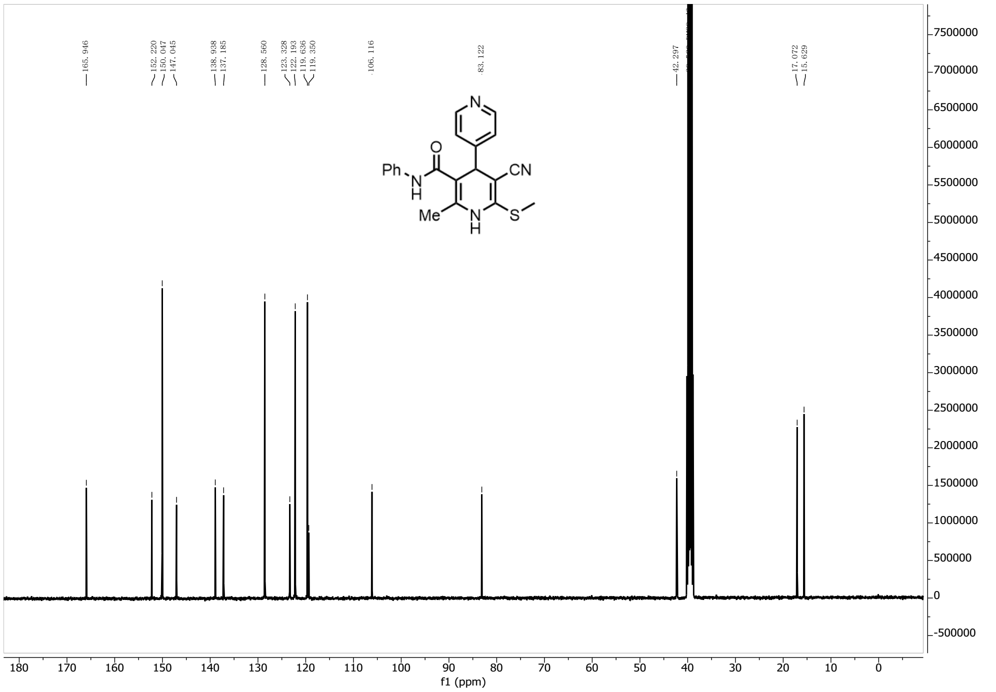


### $^{13}C$ NMR Spectrum (KL7)

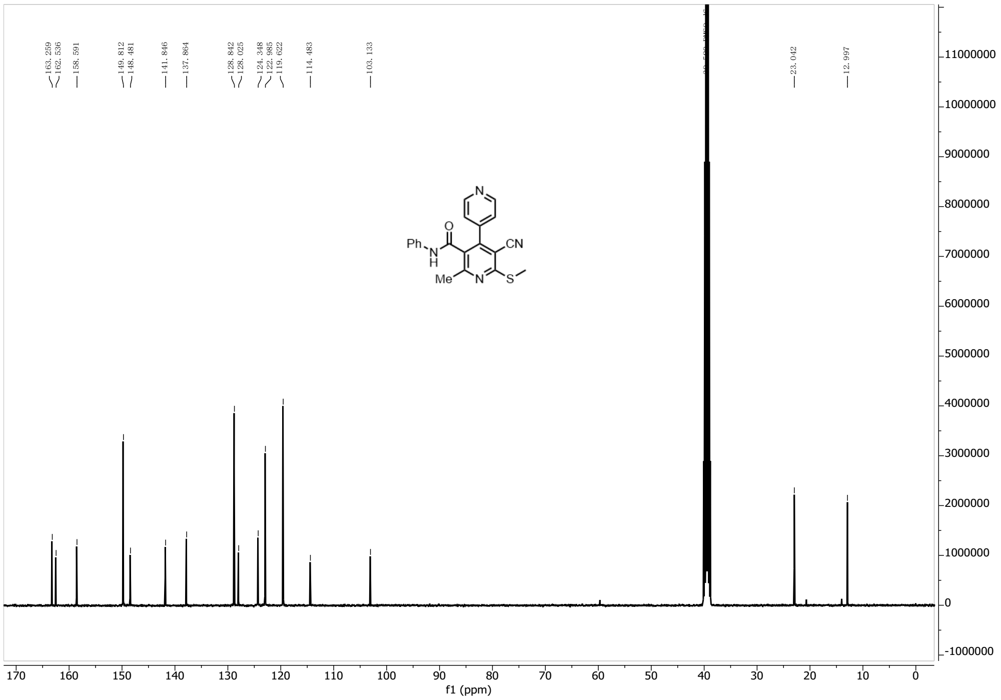


### HPLC Analysis (KL1)

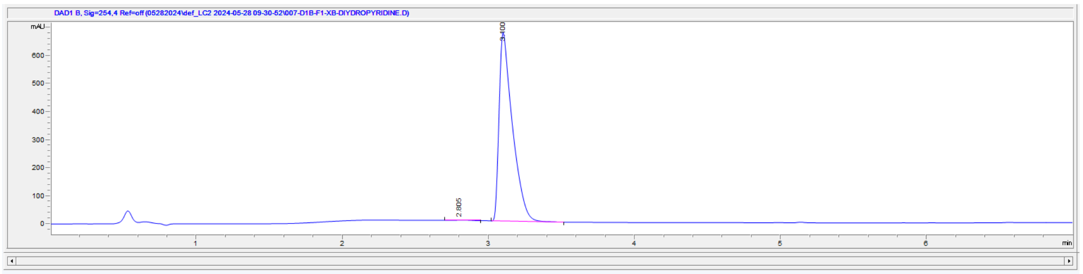


**File Information**

| | |
|---|---|
| Barcode | |
| Operator | SYSTEM |
| Method | A_LCMS_DEFAULT_METHOD_POS_CH3CN.M |
| Reference | |
| Analysis Time | 6.993 min |
| Sampling Rate | 0.0067 min (0.402 sec), 1050 datapoints |

| # | Time | Type | Area | Height | Width | Area% | Symmetry |
|---|---|---|---|---|---|---|---|
| 1 | 2.805 | BB | 10.9 | 1.3 | 0.1132 | 0.255 | 0.487 |
| 2 | 3.1 | BB | 4267 | 673.6 | 0.0919 | 99.745 | 0.404 |

### HPLC Analysis (KL7)

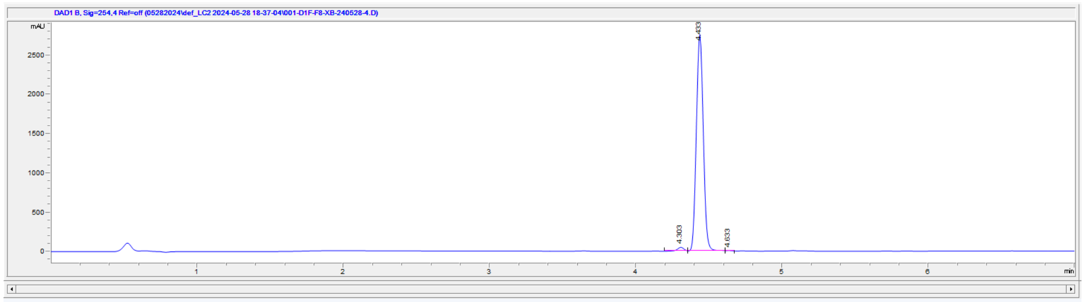


**File Information**

| | |
|---|---|
| Operator | SYSTEM |
| Method | A_LCMS_DEFAULT_METHOD_POS_CH3CN.M |
| Reference | |
| Analysis Time | 7 min |
| Sampling Rate | 0.0067 min (0.402 sec), 1051 datapoints |

| # | Time | Type | Area | Height | Width | Area% | Symmetry |
|---|---|---|---|---|---|---|---|
| 1 | 4.303 | BB | 141.5 | 46.1 | 0.0478 | 1.530 | 1.153 |
| 2 | 4.433 | BB | 9099.7 | 2766.7 | 0.0524 | 98.414 | 0.917 |
| 3 | 4.633 | BB | 5.1 | 2.3 | 0.036 | 0.056 | 0.396 |

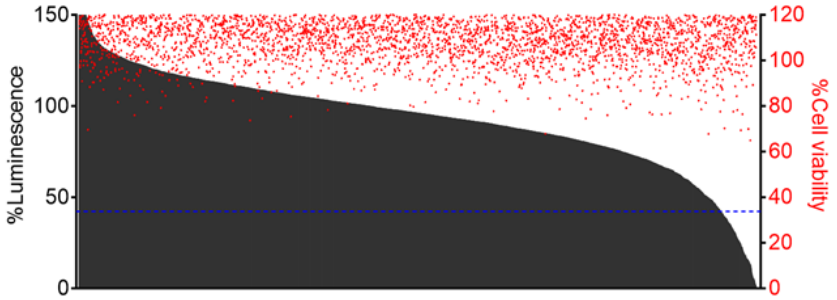


**Supplementary figure 1. High-throughput screening of a kinase-targeted library identifies compounds with differential capacities in modulating the energy state of intracellular *S. aureus*.** A total of >4,700 compounds (10 μM) from the UNC CICBDD were screened for targeting the intracellular MRSA. All of these compounds shared structural similarity to kinase inhibitors and complied with the Lipinski's rule of five. The luminescence signals were normalized to the vehicle controls (0.1% DMSO) (black bars). The normalized host cell viability was measured by the CellTiter-Fluor assay (red circles). The %luminescence of 10 μg/mL rifampicin-treated cells is indicated as a reference (blue dashed line).

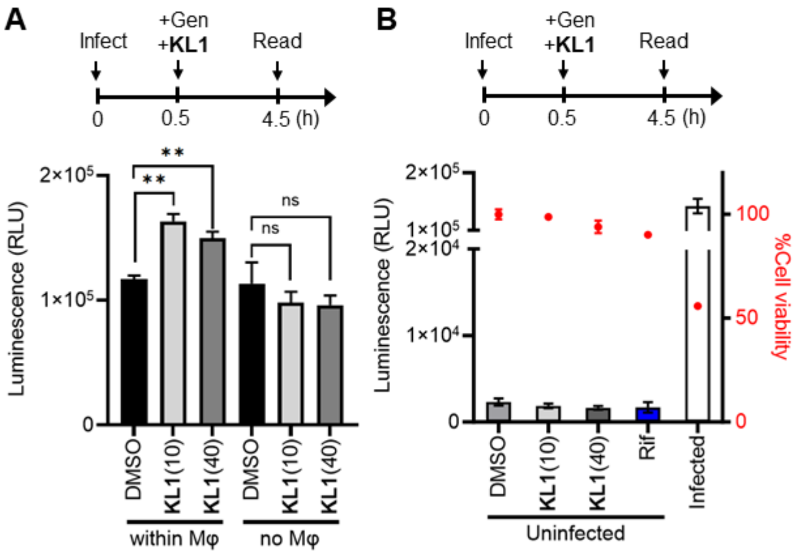


**Supplementary figure 2. The energy state up-regulation by KL1 is not caused by background noise from the compound, the host cells or dead extracellular bacteria.** (**A**) Macrophages infected with the bioluminescent MRSA strain JE2–lux (within Mφ) were treated with 10 μM or 40 μM **KL1**. Gentamicin (Gen; 50 μg/mL) was added to eliminate the extracellular bacteria. The bacterial cultures without macrophages (no Mφ) in the presence of Gen were examined with the same input CFU and culture conditions (n = 3). (**B**) Uninfected macrophages were treated with 10 μM or 40 μM **KL1**. The normalized host cell viability was measured by the CellTiter-Fluor assay (red circles). Rifampicin (Rif)-treated cells (blue bar) and infected macrophages (white bar) were included as controls (n = 3). Assay schematics are shown above plots. **p<0.01; ns, not significant (unpaired t-test). The bars represent mean ± SEM.

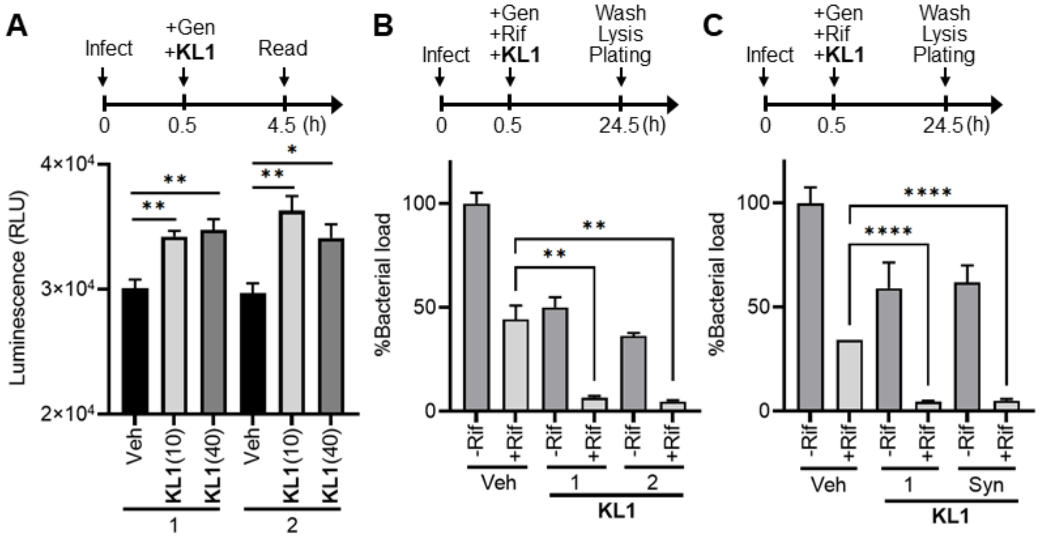


**Supplementary figure 3. The adjuvant activity of KL1 was consistently observed with the compound across different sources.** (**A** and **B**) **KL1** (10 and 40 μM) purchased from two independent suppliers, ChemBridge (1) and Enamine (2), consistently elevated the energy state of intracellular *S. aureus* (n = 4) and sensitized the bacteria to killing by rifampicin (Rif, 10 μg/mL) (n = 3). (**C**) **KL1** synthesized in-house (Syn) displayed adjuvant activity comparable to that of commercially available **KL1** (n = 3). Gentamicin (Gen; 50 μg/mL) was added to eradicate the extracellular bacteria. The numbers of tolerant bacteria were normalized to the untreated control (no Rif, no **KL1**, Gen-only). Assay schematics are shown above plots. *p<0.05; **p<0.01; ****p<0.0001 (unpaired t-test). The bars represent mean ± SEM.

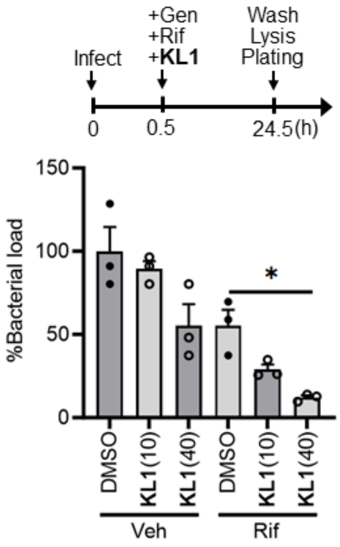


**Supplementary figure 4. Compound KL1 sensitizes the killing of intracellular MRSA by antibiotics in a dose-dependent manner.** RAW 264.7 cells were infected with the MRSA strain JE2–lux and treated with 10 μM or 40 μM **KL1** in the presence or absence of 10 μg/mL rifampicin (Rif). Gentamicin (Gen; 50 μg/mL) was added to eliminate the extracellular bacteria. The numbers of survivor bacteria were normalized to the untreated control (no Rif, no **KL1**, Gen-only) (n = 3). *p<0.05 (unpaired t-test). The bars represent mean ± SEM.

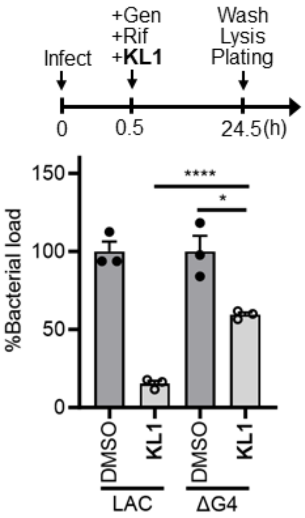


**Supplementary figure 5. Glucose uptake is important for the synergism between KL1 and antibiotics.** RAW 264.7 cells were infected with the wild-type strain LAC or the mutant lacking all four glucose transporters (ΔG4). The infected cells were treated with 40 μM **KL1** and 10 μg/mL rifampicin (Rif). Gentamicin (Gen) was included to eliminate the extracellular bacteria. The numbers of survivor bacteria were normalized to the corresponding controls (no **KL1**, Gen- and Rif-only) (black circles) (n = 3). *$p<0.05$; ****$p<0.0001$ (unpaired t-test). The bars represent mean ± SEM.

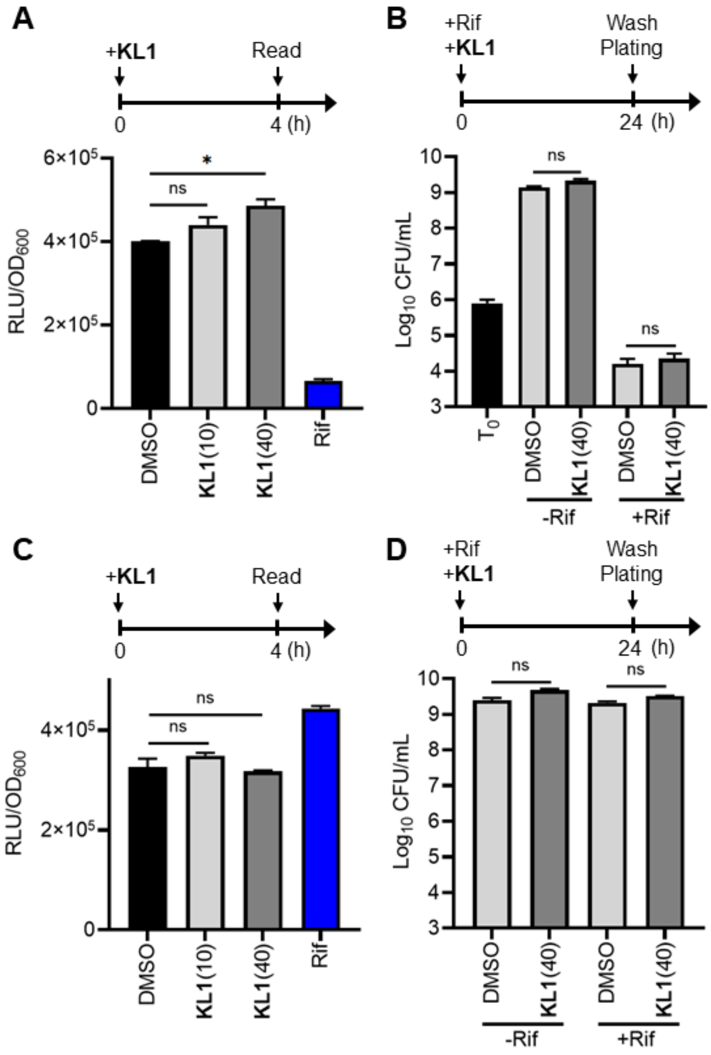


**Supplementary figure 6. Compound KL1 exhibits a minute effect on the extracellular *S. aureus* activity and does not synergize with antibiotics in killing the extracellular bacteria.** (**A** and **C**) The exponential-phase (**A**) and stationary-phase (**C**) *S. aureus* cultures (JE2–lux) were incubated with 10 μM or 40 μM **KL1** for 4 h, followed by luminescence detection. The signal intensities (RLU) were normalized to the $OD_{600}$ values. The vehicle control (0.1% DMSO, black bar) and rifampicin (Rif; 2 μg/mL)-treated group (blue bar) were included as references. (**B** and **D**) The exponential-phase (**B**) and stationary-phase (**D**) cultures were treated with and without

40 µM **KL1** and 2 µg/mL Rif for 24 h, followed by a wash step and plating to enumerate the number of tolerant bacteria. Assay schematics are shown above plots. *$p<0.05$; ns, not significant (unpaired t-test). The bars represent mean ± SEM.

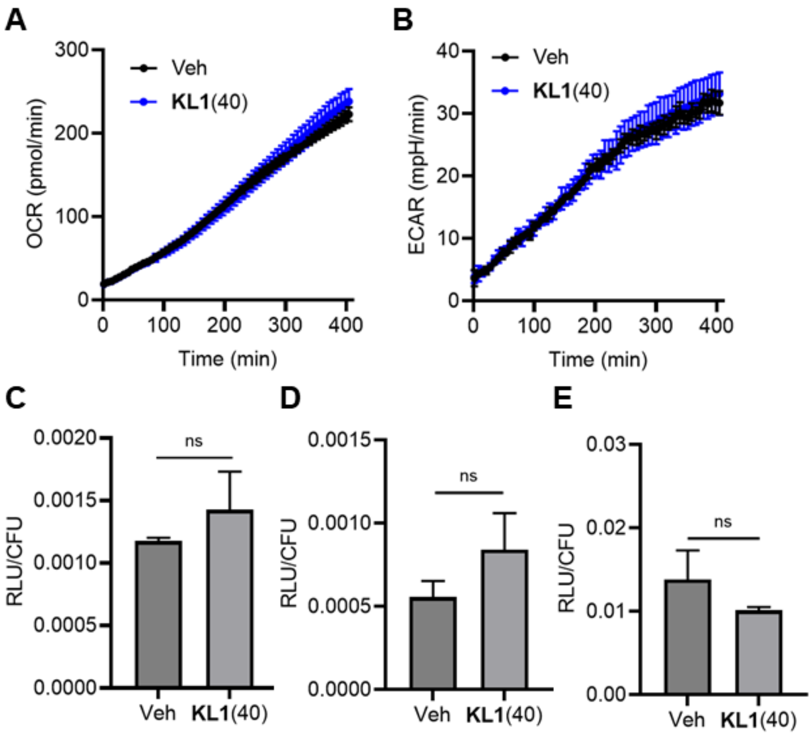


**Supplementary figure 7. Compound KL1 does not impact the metabolic activity and ATP levels in extracellular bacteria.** (**A** and **B**) Metabolic analysis indicates that **KL1** (40 μM, blue) does not alter the oxygen consumption rate (OCR) (**A**) and extracellular acidification rate (ECAR) (**B**) in extracellular *S. aureus* comparing to the vehicle control (Veh, 0.1–0.5% DMSO, black). Representative data of six independent experiments (n = 3) is shown. The bars represent mean ± SD. (**C**–**E**) Exponential-phase cultures were treated with 40 μM **KL1** or 0.1% DMSO for 4 h, followed by measuring the relative ATP levels using a BacTiter-Glo assay. The luminescent signals were normalized to the numbers of bacteria. *S. aureus* strains HG003 (**C**), JE2 (**D**) and LAC (**E**) were examined (n = 3–6). ns, not significant (unpaired t-test).

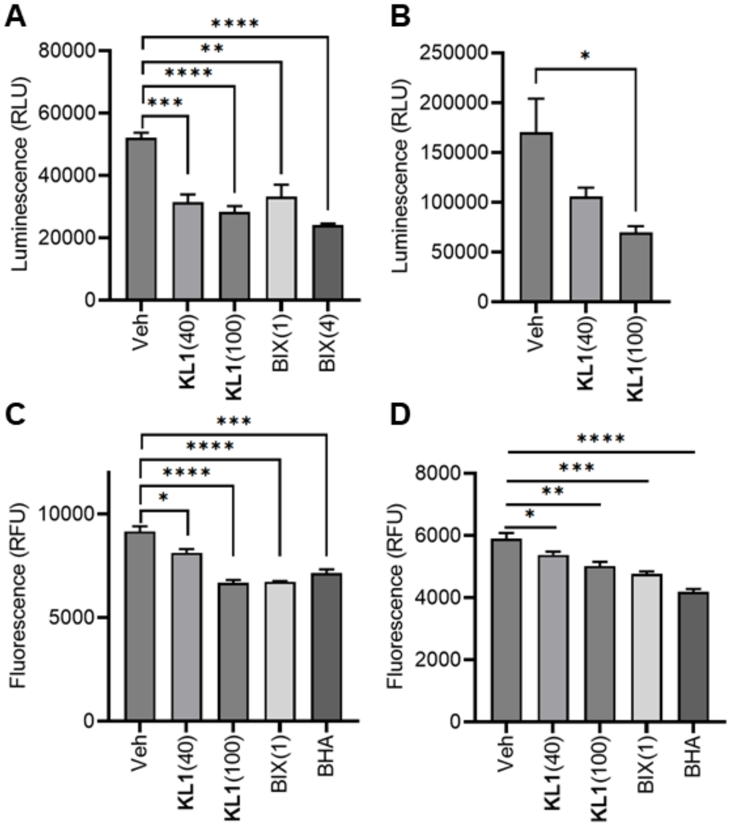


**Supplementary figure 8. KL1 reduces the level of reactive species in both infected and uninfected macrophages.** Compound **KL1** (40 and 100 µM) and the EHMT2/G9a inhibitor, BIX-01294 (BIX; 1 and 4 µM) decreased the ROS/RNS level in *S. aureus*-infected macrophages at 8 hours post-infection (hpi) (**A** and **C**) and in uninfected macrophages (**B** and **D**). L-012 (**A** and **B**) and fluorescein-boronate (Fl-B) (**C** and **D**) were used to quantify the ROS/RNS level. DMSO (Veh; 0.25%) and butylated hydroxyanisole (BHA; 20 µM) were included as controls. Representative data of three independent experiments are shown (n = 5). *p<0.05; **p<0.01; ***p<0.001; ****p<0.0001 (unpaired t-test). The bars represent mean ± SEM.

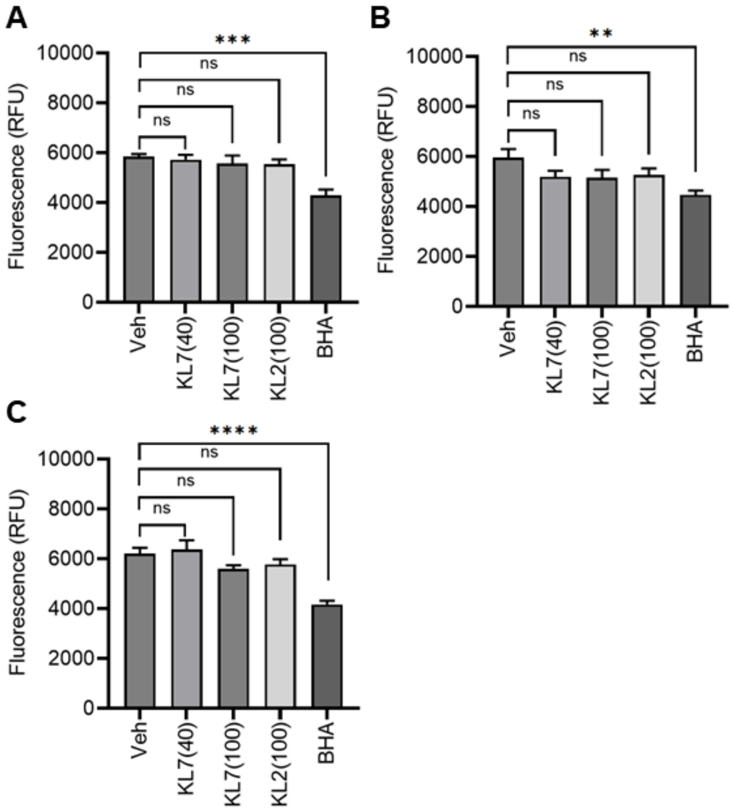


**Supplementary figure 9. The inactive analog of KL1 losses the ability to reduce the level of reactive species in both infected and uninfected macrophages.** The inactive analog KL7 (40 and 100 µM) and analog KL2 (100 µM) with lower adjuvant activity did not affect ROS/RNS production in uninfected (**A** and **B**) and *S. aureus*-infected macrophages (**C**) after 4 h (**A**) and 8 h (**B** and **C**) treatment. Fluorescein-boronate (Fl-B) was used to quantify the level of reactive species. DMSO (Veh; 0.25%) and butylated hydroxyanisole (BHA; 20 µM) were included as controls. Representative data of three independent experiments are shown (n = 5). **p<0.01; ***p<0.001; ****p<0.0001; ns, not significant (unpaired t-test). The bars represent mean ± SEM.

**Supplementary table 1. Primers for molecular cloning.**

| Primer | Sequence |
|---|---|
| mKate forward | CAAATAGGTACCTATGTCAGAACTTATCAAGGAAAATATG[†] |
| mKate reverse | GATTACGAATTCTTAACGGTGTC |
| mKate_597[‡] | CTTGTCGGTGGAGGTCAC |
| mKate CTC forward | CCTTGATAAGTTCTGACATAGGTACCATCctcCTTATTTTAATTATACTCTATCAATGATAG[§] |
| mKate GAG reverse | CTATCATTGATAGAGTATAATTAAAATAAGgagGATGGTACCTATGTCAGAACTTATCAAGG[§] |

[†]The sequence in red represents 5' overhang containing a KpnI site.

[‡]Sequencing primer to confirm the identity of the gene.

[§]The sequence in lowercase represent inserted nucleotides for incorporating an upstream ribosomal binding site.

**Supplementary table 2. Minimum inhibitory concentrations (MICs) of antibiotics in the *S. aureus* strains used in this study.**

| Antibiotics | LAC | JE2–Lux | High persister | Low persister |
|---|---|---|---|---|
| Rifampicin | 0.008±0.004 μg/mL | 0.006 μg/mL | 0.006 μg/mL | 0.008±0.004 μg/mL |
| Moxifloxacin | 5 μg/mL | 5 μg/mL | 0.16 μg/mL | 0.16 μg/mL |
| Vancomycin | 1.33±0.58 μg/mL | 1 μg/mL | 1.33±0.58 μg/mL | 1 μg/mL |

Three independent experiments were conducted, and the MIC values are presented as the average ± the standard deviation.

**Supplementary table 3. Potential targets of KL1 based on published functional screens.**

| Target | BioAssay ID[†] | Assay | Activity | Location, expression[‡] |
|---|---|---|---|---|
| PHOSPHO1 | 1565 | uHTS absorbance assay for the identification of compounds that inhibit PHOSPHO1 | 61.5% inhibition at 13.3 μM | Intracellular, immune cells and other cell types |
| SLC5A7 | 488975 | Primary cell-based screen for identification of compounds that inhibit the Choline Transporter | BScore_intRatio of -5.0281 | Membrane, not detected in immune cells |
| SLC5A7 | 493221 | Confirmatory screen for compounds that inhibit the Choline Transporter | 50.55% inhibition at 10 μM | Membrane, not detected in immune cells |
| EHMT2/G9a | 504332 | qHTS Assay for Inhibitors of Histone Lysine Methyltransferase G9a | $EC_{50}$ value of 25.12 μM | Intracellular, immune cells and other cell types |
| COPS5 | 651999 | uHTS identification of small molecule inhibitors of Csn-mediated Deneddylation of Cullin-Ring Ligases, vis a fluorescence polarization assay | 93.48% inhibition at 12.5 μM | Intracellular, immune cells and other cell types |

[†]Data sourced form PubChem biological test results.

[‡]Data sourced form Expression Atlas.

**Movie S1.** Confocal z-sections of the representative macrophages infected with an inducible GFP reporter *S. aureus* strain (green).

**Movie S2.** Live imaging of representative macrophages infected with an inducible mKate reporter *S. aureus* strain (red). Cells were stained with Hoechst 33342 (blue) and LysoTracker DND-26 (green) to visualize the nucleus and lysosomes.

**Movie S3.** Confocal z-sections of representative *S. aureus* (red)-infected macrophages. Cells were stained with Hoechst 33342 (blue) and LysoTracker DND-26 (green) to visualize the nucleus and lysosomes.
